# SARS-CoV-2 infects human adipose tissue and elicits an inflammatory response consistent with severe COVID-19

**DOI:** 10.1101/2021.10.24.465626

**Authors:** Giovanny J. Martínez-Colón, Kalani Ratnasiri, Heping Chen, Sizun Jiang, Elizabeth Zanley, Arjun Rustagi, Renu Verma, Han Chen, Jason R. Andrews, Kirsten D. Mertz, Alexandar Tzankov, Dan Azagury, Jack Boyd, Garry P. Nolan, Christian M. Schürch, Matthias S. Matter, Catherine A. Blish, Tracey L. McLaughlin

## Abstract

The COVID-19 pandemic, caused by the viral pathogen SARS-CoV-2, has taken the lives of millions of individuals around the world. Obesity is associated with adverse COVID-19 outcomes, but the underlying mechanism is unknown. In this report, we demonstrate that human adipose tissue from multiple depots is permissive to SARS-CoV-2 infection and that infection elicits an inflammatory response, including the secretion of known inflammatory mediators of severe COVID-19. We identify two cellular targets of SARS-CoV-2 infection in adipose tissue: mature adipocytes and adipose tissue macrophages. Adipose tissue macrophage infection is largely restricted to a highly inflammatory subpopulation of macrophages, present at baseline, that is further activated in response to SARS-CoV-2 infection. Preadipocytes, while not infected, adopt a proinflammatory phenotype. We further demonstrate that SARS-CoV-2 RNA is detectable in adipocytes in COVID-19 autopsy cases and is associated with an inflammatory infiltrate. Collectively, our findings indicate that adipose tissue supports SARS-CoV-2 infection and pathogenic inflammation and may explain the link between obesity and severe COVID-19.

**One sentence summary:** Our work provides the first *in vivo* evidence of SARS-CoV-2 infection in human adipose tissue and describes the associated inflammation.

## INTRODUCTION

The COVID-19 pandemic, caused by the viral pathogen SARS-CoV-2, has infected over 250 million people and taken the lives of over 4.5 million individuals globally as of October 2021. The clinical manifestations of COVID-19 range from asymptomatic infection, mild cold-like symptoms, to severe pulmonary and extrapulmonary manifestations characterized by extreme inflammation and cytokine storm (*1*). Obesity has emerged as an independent risk factor for infection, severe disease, and mortality (*2–6*). While obesity is associated with comorbid conditions also related to severe COVID-19, the independent relative risk of obesity is higher than that of hypertension and type 2 diabetes (*2, 5, 7*). Further, obesity is a risk factor even in young adults and children who do not have other comorbid conditions (*8*). Several distinct mechanisms could underlie this association. Impaired respiratory mechanics may result from a heavy chest wall, airway resistance, and/or presence of obstructive sleep apnea (*9*). The metabolic milieu in obesity, particularly among individuals with insulin resistance, is characterized by systemic inflammation and hypercoagulability (*10–12*) and could thus stimulate a more robust inflammatory response to SARS-CoV-2. Impaired immune responses to viral infection are another possibility, as obese individuals exhibit altered immune cell profiles at baseline (*13*) and in response to influenza infection (*7, 14*).

Furthermore, a recent report demonstrated that SARS-CoV-2 can infect adipocytes *in vitro* (*15*); it is not known whether SARS-CoV-2 infects other adipose tissue-resident cells and/or drives an inflammatory response in adipose tissue. Other viruses have been shown to infect several cell types within adipose tissue, including adipocytes (influenza A virus), adipose-stromal cells (adenovirus 36, human cytomegalovirus), macrophages (simian immunodeficiency virus (SIV)), and T cells (human immunodeficiency virus (HIV))(*16–19*). Complex interactions between various cell types and adipocytes can drive significant inflammation, with reports of adipocyte-derived chemoattractants such as monocyte chemoattractant protein-1 (MCP-1) leading to macrophage infiltration (*20*), tumor necrosis factor alpha (TNF-ɑ) activating nuclear factor kappa B (*21, 22*), and free fatty acids driving toll-like receptor 4-mediated inflammation and insulin resistance (*23*). Thus, pronounced and/or prolonged SARS-CoV-2 viral replication and inflammation might occur in those with obesity and contribute to severe disease. Of particular concern is the possibility that viral infection of peri-organ fat could contribute to organ damage via inflammation and downstream processes such as extracellular matrix deposition/fibrosis, edema, impaired cellular function, endothelial dysfunction, and hypercoagulability (*11, 13*). To date, COVID-19 profiling studies have generally excluded adipose tissue from analyses; in other cases adipose tissue may have been lumped in with analyses of adjoining tissues (*24, 25*) These studies have shown that SARS-CoV-2 RNA and proteins are detected across numerous tissues, including the lung, brain, intestine, and pancreas (*24, 26, 27*). Thus, we undertook a study to test the hypothesis that SARS-CoV-2 infects cells within human adipose tissue and incites an inflammatory response. We harvested adipose tissue from multiple depots in uninfected obese humans for *in vitro* infection and obtained autopsy specimens of various adipose depots in individuals who died from COVID-19. Our results clearly show SARS-CoV-2 infection in macrophages and adipocytes from multiple adipose depots, with an attendant increase in inflammatory profile.

## RESULTS

### Establishment of a system to examine SARS-CoV-2 infection in human adipose tissue

To explore the ability of human adipose tissue to support SARS-CoV-2 infection and drive an inflammatory response, we recruited participants undergoing bariatric or cardiothoracic surgery. Freshly harvested tissue from the subcutaneous (SAT), visceral (VAT), pericardial (PAT) and epicardial (EAT) adipose tissue depots was subjected to collagenase digestion to separate stromal-vascular cells (SVC) from mature adipocytes. Clinical characteristics are shown in table S1.

### Human adipose tissue SVC supports SARS-CoV-2 infection

Freshly isolated SVC from human adipose tissue were exposed to SARS-CoV-2 (WA-01) at a multiplicity of infection (MOI) of 1 or mock-infected. Infection and replication were assessed by reverse transcription quantitative PCR (RTqPCR) for SARS-CoV-2 nucleocapsid (*N*) gene and by flow cytometry for SARS-CoV-2 N protein (Figure 1A). Both genomic (Fig. 1B) and subgenomic (fig. S1) viral RNA were detected in SARS-CoV-2 infected SVC from subcutaneous (n=6 Fig.1B; n=2 fig. S1), omental (n=4 Fig.1B; n=2 fig. S1), pericardial (n=1 fig. S1), and epicardial (n=1 fig. S1) adipose tissue. To confirm that *N-gene* detection represented active viral infection, we cultured infected cells in the presence of remdesivir, an inhibitor of the viral RNA-dependent RNA polymerase. We first confirmed the ability of remdesivir to inhibit viral RNA accumulation in A549-ACE2 cells (fig. S2). In SARS-CoV-2-infected SVC from both SAT and VAT, we observed accumulation of *N-gene* copies over the course of 96 hours, indicating viral RNA replication (Fig. 1B). Importantly, remdesivir significantly reduced viral RNA levels at later time points in both tissue types (Fig. 1C). Together, these data indicate that SVC can support SARS-CoV-2 infection.

**Figure 1.**
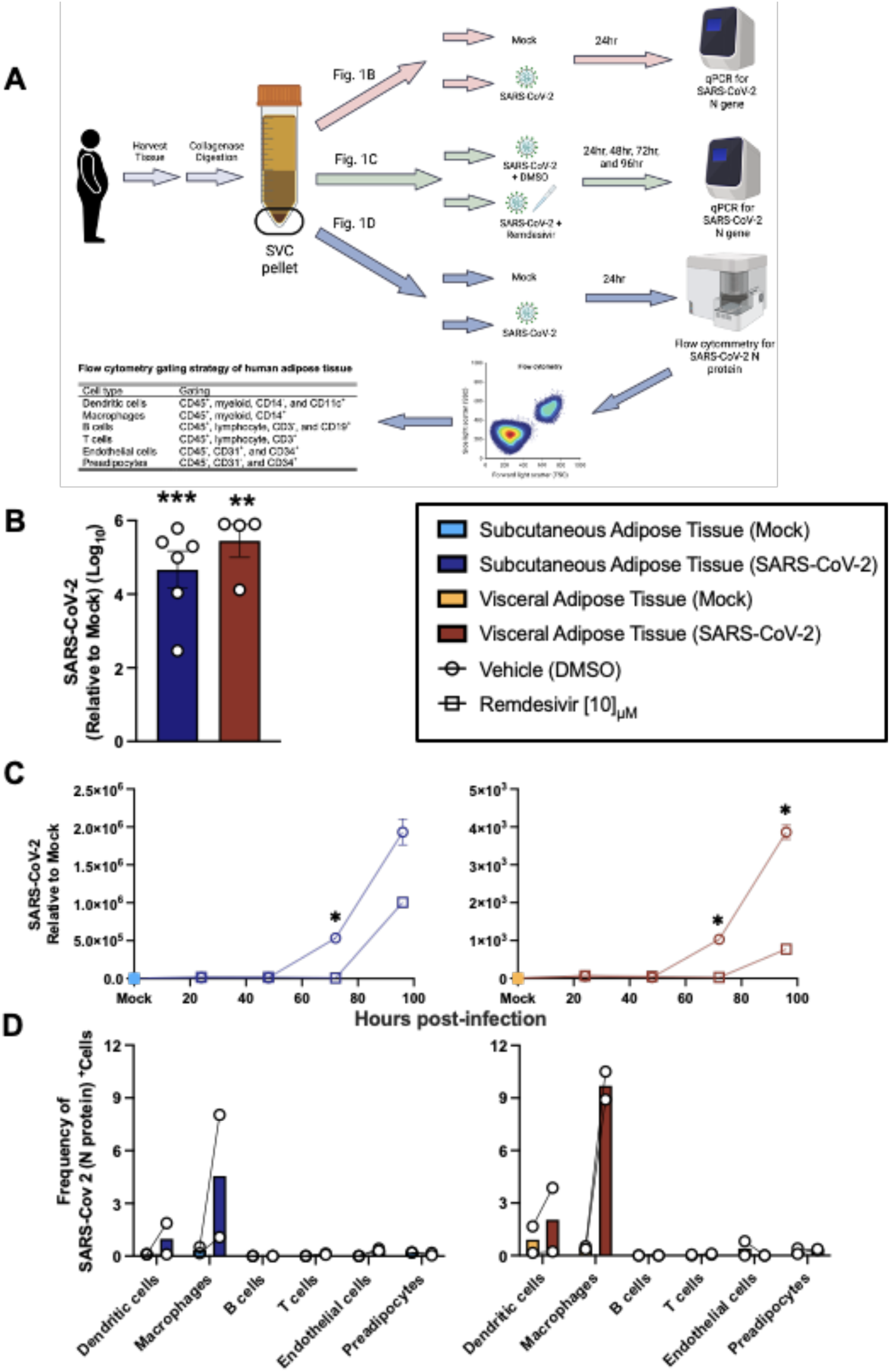
Exposure of stromal vascular cells (SVC) from adipose tissue to SARS-CoV-2 supports infection of macrophages. (A) Sketch of workflow. SVC of human adipose tissue was isolated by collagenase digestion prior to viral infection. SVC was infected or left untreated (mock) with SARS-CoV-2 (USA-WA1/2020) at a multiplicity of infection (MOI) of 1. (B) Relative gene expression of SARS-CoV-2 (N gene) obtained by 1-step RTqPCR.at 24 hpi or mock infection (SAT, n=6; VAT, n=4). (C) Relative gene expression of SARS-CoV-2 (N gene) obtained by 1-step RTqPCR in cultures were treated with vehicle (DMSO) or 10µM remdesivir to inhibit viral replication and maintained for 24, 48, 72, and 96 hours before harvest. Relative expression was analyzed by ΔΔCt method relative to a mock sample using 18S rRNA as a housekeeping gene. Each data point is an average of 3 technical replicates. (D) Frequency of SARS-CoV-2 infected cells based on SARS-CoV-2 N protein detection by flow cytometry of SAT (left, n=2) and VAT (right, n=2). Gating is detailed in supplemental figure 3. Statistical analysis: (A) paired, two-sided, student’s t-test. **P<0.01, ***P<0.001. (B) Statistical analysis was performed with a two-way ANOVA, multiple comparisons using statistical hypothesis Sidak. *P<0.05. Data is presented as ± mean s.e.m. (A) Sketch was created with BioRender.

Next, to determine which of the SVC are infected, we performed flow cytometry to assess viral N protein expression in six major SVC types (dendritic cells, macrophages, B cells, T cells, preadipocytes, and endothelial cells) (Fig. 1A and fig. S3). In both SAT and VAT, SARS-CoV-2 N protein was primarily restricted to macrophages (defined as large granular CD45^+^CD14^+^CD11c^-^ cells) (Fig.1D). We next assessed the level of angiotensin-converting enzyme 2 (ACE-2), the cellular receptor for SARS-CoV-2 (*28*) in the SVC. ACE-2 protein expression was very limited in SVC, with no detectable expression in the VAT and very low-level expression on ∼3% of SAT macrophages when compared to isotype control (fig. S4). In addition, we did not detect *ACE2* mRNA in any of the six samples collected from SAT, VAT, PAT or EAT (table S2). Thus, these data indicate that SARS-CoV-2 primarily infects macrophages in SVC and may enter through a non-canonical entry receptor.

### SARS-CoV-2 infects a distinct subset of macrophages

To comprehensively characterize the cells infected with SARS-CoV-2 in human adipose tissue, we performed single-cell RNA sequencing (scRNA-seq) of SARS-CoV-2 and mock-infected SVC isolated from both SAT and VAT from three bariatric surgery patients (Fig. 2A, table S3). We generated 198,759 single-cell expression profiles. After performing dimensionality reduction and Harmony batch integration (*29*), unbiased graph-based clustering resulted in 23 cell types (C0-C22) with marker genes that aligned with previous datasets and included three major cell groups: preadipocytes, immune cells, and endothelial cells (*30, 31*) (Fig. 2B, table S3, fig. S5). We saw the greatest heterogeneity across the preadipocyte population for which we identified 14 distinct clusters, each of which was labeled by its top two cluster defining genes. We similarly labeled two distinct macrophage clusters.

**Figure 2.**
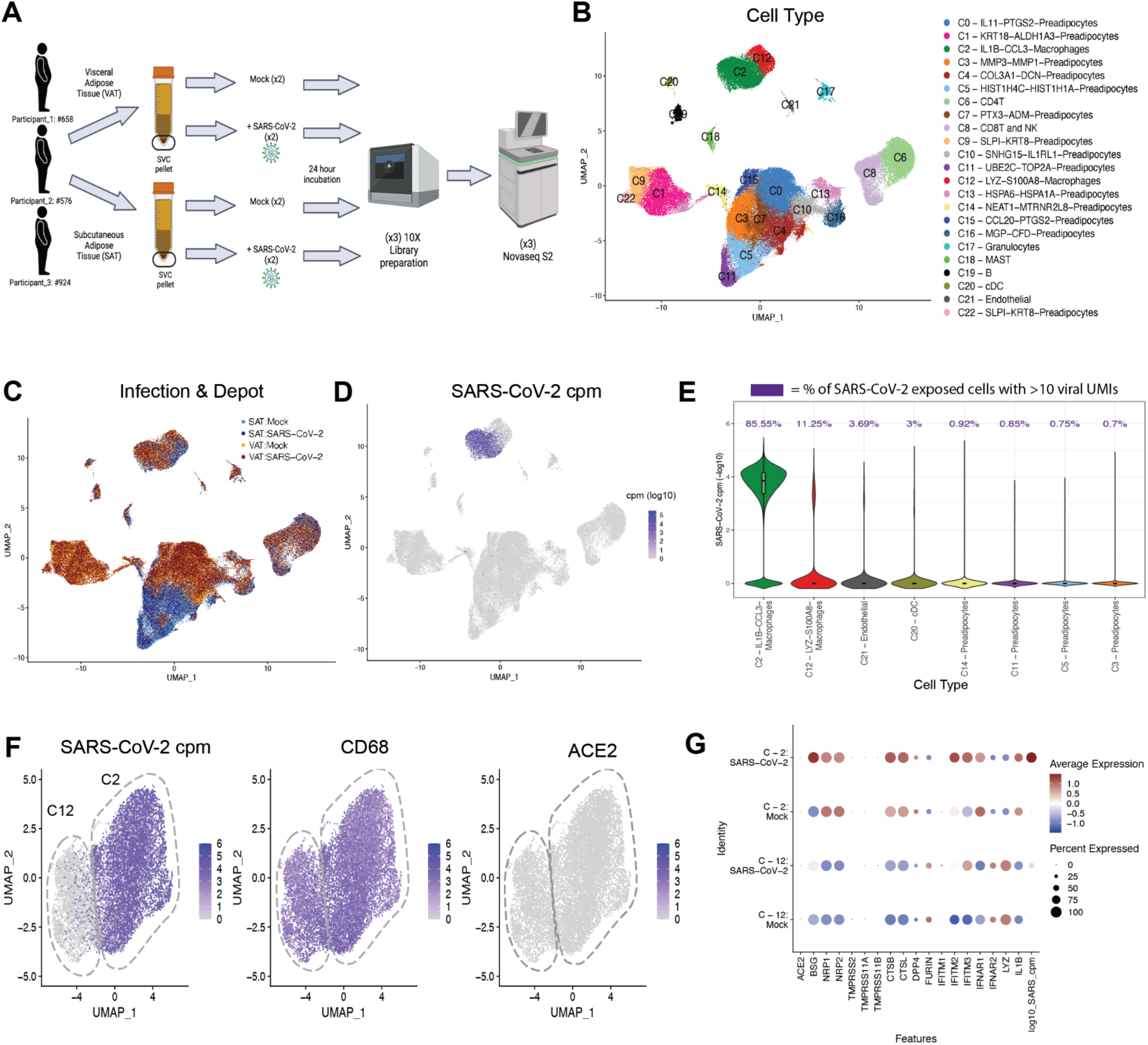
A subset of macrophages is infected with SARS-CoV-2. **(A)** Schematic of experiment. The stromal vascular cells (SVC) were isolated from the SAT and VAT depots of three different participants and infected with either mock or SARS-CoV-2 (MOI of 1.0). Each sample was collected for scRNA-seq at 24 hpi. **(B-C)** UMAP representation of the SVC from all participants (n=3) across 198,759 cells, **(B)** colored by manually annotated cell type and **(C)** colored by infection and depot. **(D)** UMAP representation of all cells colored by SARS-CoV-2 cpm (log10). **(E)** Violin plot reveals the SARS-CoV-2 cpm values of all cells across SARS-CoV-2 infected samples, showing only the 8 cell clusters with the highest composition of SARS-CoV-2^+^ cells. Percentages above each cell type denote the percentage of cells within each cluster that have over 10 SARS-CoV-2 reads. **(F)** UMAP projections of all macrophages from the scRNA- seq dataset, colored by SARS-CoV-2 cpm (left), colored by CD68 expression (middle) and colored by ACE2 expression (right). **(G)** Dotplot of the proportion of cells (dot size) in the macrophage clusters split by infection condition expressing genes relevant for SARS-CoV-2 entry and antiviral defense, as well as SARS-CoV-2 cpm and macrophage cluster markers and colored by scaled average expression. (A) Figure was created with BioRender.

We detected SARS-CoV-2 at levels ranging up to 1079 transcripts per cell in a total of 8.8% of cells in the infected samples (fig. S6) and did not detect a single SARS-CoV-2 transcript in the mock-infected samples. The uniform manifold approximation and projection (UMAP) of SARS-Cov-2-exposed samples colored by viral counts-per-million (cpm) showed that macrophages contained the highest concentration of SARS-CoV-2 transcripts (Fig. 2D), with >85% of SARS-CoV-2 exposed C2-macrophages and >11% C12-macrophages containing more than ten viral transcripts (Fig. 2E). *ACE2* transcript was not detected in the SVC macrophages and therefore could not explain the difference in infection levels between C2-macrophages and C12-macrophages (Fig. 2F, 2G). However, C2-macrophages expressed higher levels of *BSG*, *NRP1/2*, *CTSB* and *CTSL* compared to C12-macrophages (Fig. 2G), all previously noted to be important to SARS-CoV-2 infection and viral processing (*32–34*). Our data suggest that these alternative entry receptors could contribute to SARS-CoV-2 entry into C2-macrophages.

To further evaluate how these two macrophage populations differ in SARS-CoV-2 susceptibility, we examined the most differentially expressed genes between the macrophage cluster that was highly enriched for SARS-CoV-2 reads (C2) versus the relatively uninfected cluster (C12) at baseline (mock-infected) and identified 1000 significant differentially expressed genes (Fig 3A, table S4). Notably, even in the mock condition, there was a significant upregulation of inflammation related transcripts within C2 compared to C12, and in particular, transcripts for numerous chemokine ligands (Fig. 3A, fig. S7B, table S6-7). Increased expression of *CCL8* and *CCL3*, chemokines involved in monocyte chemotaxis, highlight the potential of these infected cells to induce the infiltration of inflammatory monocytes and macrophages (*35, 36*). There was significant downregulation of 14 HLA genes across HLA Class I and Class II proteins, including *HLA-DRB1*, *HLA-DRA*, *HLA-DQB1*, and *HLA-DPB1*, in the C2-macrophages compared to C12-macrophages (Fig. 3A, table S4). Additionally, the C2-macrophage cluster demonstrated enrichment for transcripts associated with perivascular macrophages (PVM) including *CD163*, *SELENOP*, *MARCO* and *APOE* (Fig. 3B and (*31, 37, 38*)). In contrast, the relatively uninfected C12 demonstrated upregulation of genes relating to lipid metabolism and other markers of lipid-associated macrophages, including *FABP4*, *TGM2*, *GSN, TREM2, LPL* and *PGD,* suggesting that this macrophage population might be implicated in lipid accumulation and trafficking and derived from metabolic activation (*30, 31, 37, 39*).

**Figure 3.**
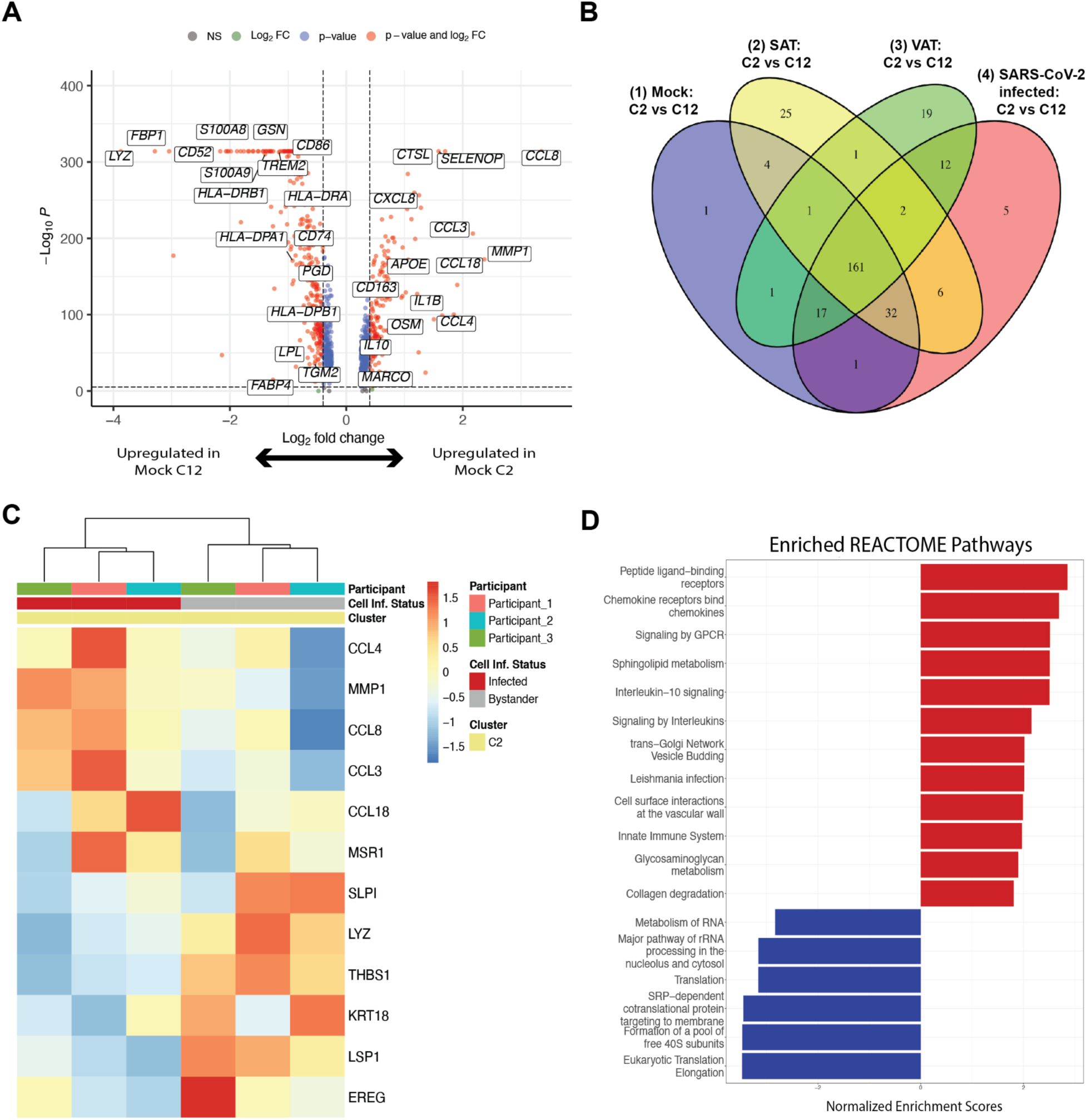
The infected macrophage cluster is marked by increased chemokine expression. **(A)** Volcano plot of the differentially expressed genes between macrophage clusters 2 (C2) and 12 (C12) across mock-infected samples. **(B)** Venn Diagram comparison of the significantly differentially expressed genes (DEGs) and their direction of change across C2 versus C12 in (1) mock-infected, (2) all SAT and (3) all VAT, and (4) SARS-CoV-2-infected conditions. **(C)** Heatmap of the most significant DEGs between SARS-CoV-2+ versus bystander macrophages within C2. **(D)** Normalized enrichment scores of top Reactome pathways, using significant DEGs between SARS-CoV-2+ versus bystander C2 macrophages.

### SARS-CoV-2 infection drives an inflammatory response in macrophages

As major differences between the macrophage clusters include markers of inflammation (i.e. *IL1B, CXCL8, CCLs*) (Fig. 3A-B), this raises the possibility that *in vitro* SARS-CoV-2 infection drives the formation of the C2-macrophage cluster. However, this cluster is present in mock-infected samples, and both SARS-CoV-2-infected and mock-infected samples displayed a similar array of differentially expressed genes between these clusters (Fig. 3B, fig. S7A), indicating that this inflammatory C2-macrophage cluster is naturally present in the SVC of adipose tissue and is more susceptible to infection *in vitro*.

To explore the cell-intrinsic effects of infection, we compared the transcriptional profiles of C2- macrophages that did (infected) or did not (bystander) contain SARS-CoV-2 transcripts within the infected cultures (fig. S7C, Fig. 3C, table S5). Though there were baseline differences in gene expression between participants, several genes strongly distinguished SARS-CoV-2^+^ C2 cells from bystander C2 cells by hierarchical clustering (Fig. 3C); many of these genes also distinguish C2-macrophages from C12-macrophages (fig. S7D). For instance, *CCL4, CCL8,* and *CCL3* genes were significantly enriched in the infected cells within C2, and *LYZ* and *THBS1* were significantly downregulated within the infected cells. As these genes contributed to defining the C2 population from C12 cells, this result suggests that SARS-CoV-2 infection may drive C2 features to further extremes. Pathway analysis of significant DEGs between the SARS-CoV-2^+^ cells versus bystanders highlights the enrichment of pathways associated with IL-10 signaling (Fig. 3D, table S8), which is consistent with upregulation of this pathway in monocytes of severe COVID-19 patients (*40*). Chemokine and other innate immune system-associated pathways are also highly enriched in infected cells (Fig. 3D), which has been previously demonstrated (*41*). Interestingly, there is a reduced enrichment in pathways related to translation machinery of the host, which is supported by SARS-CoV-2 studies suggesting that SARS-CoV-2 infection has the ability to reshape translation, splicing, protein homeostasis and nucleic acid metabolism pathways (*42, 43*). Thus, the macrophages with detectable SARS-CoV-2 RNA display a dramatic transcriptional response with upregulation of inflammatory pathways.

### Adipocyte progenitors demonstrate an inflammatory response to SARS-CoV-2 infection of macrophages

Given that exposure of preadipocytes to inflammatory cytokines such as TNF-α or IL-6 can alter their phenotype (*31, 44*), we next evaluated whether SARS-CoV-2 infection of SVC led to a bystander activation of preadipocytes. We embedded only preadipocytes and used unsupervised clustering to identify 17 unique clusters (P0-P16) (Fig. 4A-C, fig. S8A, table S9). Notably, SAT and VAT preadipocytes were highly transcriptionally unique, with 11/17 clusters each composed of over 75% of cells derived from only one depot. Overall, there was not a dramatic shift in cluster composition of SARS-CoV-2-infected and mock-infected samples, except for cluster P15 which was highly enriched with preadipocytes from SARS-CoV-2-infected SAT (Fig. 4C). This cluster was highly inflammatory, with dramatic upregulation of two antiviral interferon stimulated genes (ISGs), *IFIT1* and *ISG15* (fig. S8A), suggesting a relatively robust response to infection in the SAT. The remaining clusters were distinguished by several unique DEGs (fig. S8A, table S9), including *CXCL14* and *APOD* in cluster P9, indicating enrichment with adipocyte stem cells (ASCs) (*30, 31, 45*)*; KRT18* and *MSLN* in clusters P1 and P4 indicating enrichment with mesothelial cells in VAT only, as previously reported (*30, 45, 46*). Further, VAT-predominant clusters demonstrated higher levels of *IFI27* (a marker of SARS-CoV-2 infection in the blood) than other clusters while *IL-6* expression was high across all preadipocytes in mock-conditions, highlighting the highly inflammatory nature of preadipocytes at baseline.

**Figure 4.**
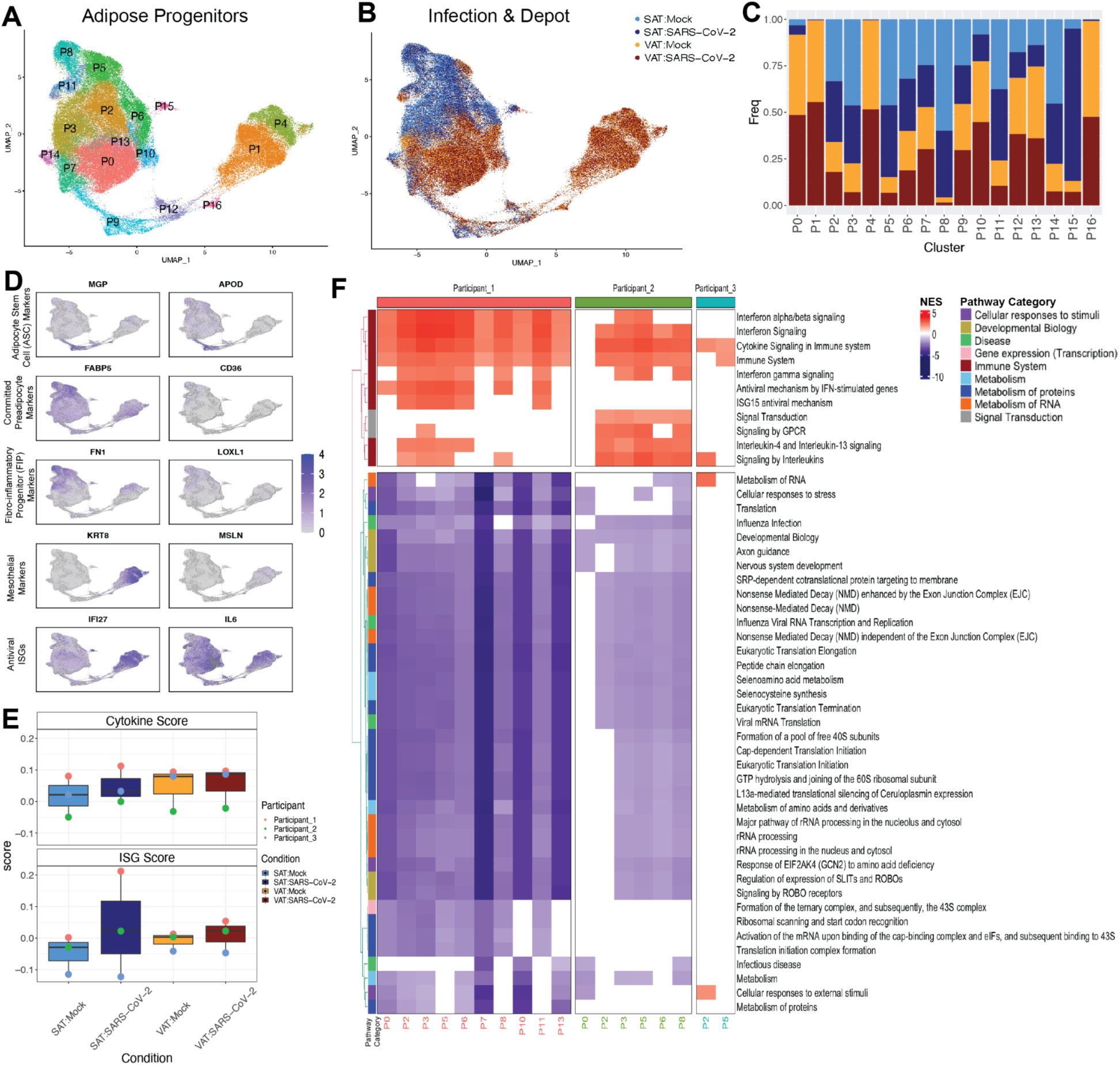
Preadipocytes respond to SARS-CoV-2 exposure. **(A, B)** UMAP embedding of all preadipocytes (n=140,867) colored by **(A)** cluster and **(B)** sample and infection type. **(C)** Cell fraction bar plot clustered by sample and infection type within each cluster. **(D)** Feature plots depicting expression of selected markers associated with preadipocyte cell states, cell types and antiviral genes. **(E)** Box plots of average cytokine (top) and ISG- (bottom) module scores across the preadipocytes of each participant and depot in both mock and SARS-CoV-2 infection conditions. **(F)** Reactome pathway analysis was performed on the significant DEGs by participant and cluster within SAT. Pathways that were represented and significant in at least four of the participant-cluster subsets were included. Pathways clustered by Euclidean distance (tree not shown) and split by the two major subtrees.

To more globally investigate the effects of SARS-CoV-2 infection on SAT and VAT preadipocytes, we looked at the total number of DEGs upon SARS-CoV-2 infection across each participant’s SAT and VAT and noticed that for each participant, the SAT had a higher number of DEGs than the paired VAT sample, suggesting that the SAT had a stronger transcriptional response than VAT (fig. S8B). We then used known ISGs and cytokine genes to generate ISG and cytokine gene scores for each participant, infection condition and depot (Fig. 4E, table S14). Infection induced both ISGs and cytokines, but SAT demonstrated a greater increase in ISG and cytokine responses than did VAT. These data also revealed that in these three participants, the VAT depot has a higher baseline cytokine expression that is further increased upon SARS-CoV-2 exposure (Fig. 4D, fig. S9A).

Next, we wanted to see whether particular preadipocyte clusters within the SAT and VAT showed a stronger response to SARS-CoV-2 exposure. We employed perturbation analysis previously described (*47, 48*) to identify which cluster within each depot and participant demonstrated the strongest changes across infection conditions (fig. S8C, S8D). Interestingly, across both depots, the ASC-like cluster, P9, along with nearby clusters, P16 and P12, showed the lowest perturbation relative to other clusters. While donors varied in their most highly perturbed clusters, this analysis demonstrated that SAT showed consistently stronger perturbation than VAT upon SARS-CoV-2 infection. Therefore, we then focused on the SAT to understand the particular pathways by cluster that contribute to the overall inflammatory response in the depot. To do this, we identified the DEGs between SARS-CoV-2 exposed versus mock preadipocytes by cluster, depot, and participant and employed gene set enrichment analysis using the Reactome database to identify relevant pathways to understand the response within each participant and each cluster (Fig. 4F, table S10-13). Clusters P5 and P2 consistently showed transcriptional enrichment for genes relevant to an immune response, more specifically related to cytokine signaling. Additionally, pathway analysis by cluster and participant shows that in participant 1 and 2, which seemed to have a more dramatic response to SARS-CoV-2 infection, we see most of the adipose clusters showing enrichment in responses relevant to the immune system. Participant 3 only had 2/17 clusters revealing enrichment for immune-related pathways. Negatively enriched pathways relate to viral transcription and translation pathways which suggests that preadipocytes are also expressing genes to suppress viral production. A similar analysis was also performed on the VAT depot which demonstrated a similar enrichment for immune response pathways across preadipocyte clusters, with participant 1 eliciting more dramatic changes across their preadipocyte compartment (fig. S8D). Overall, these data indicate that SARS-CoV-2 infection within the SVC macrophages drives inflammatory responses in the neighboring preadipocyte cells.

### SARS-CoV-2 infection of SVC induces inflammation

To better understand the impact of SARS-CoV-2-mediated infection on inflammatory responses within the SVC, we measured the relative gene expression of *IL-6* by RTqPCR (fig. S11A) in SARS-CoV-2 infected versus mock-infected SVC. We selected *IL-6* due to its high presence in severe cases of COVID-19 (*65*), its association with low-grade inflammation in obese individuals (*66*), and its likely role in COVID-19 immunopathogenesis (*67*). SARS-CoV-2-infection drove significantly increased *IL-6* expression in SAT (n=6) (fig. S11A). *IL-6* transcripts were also increased in VAT (n=6), but the change was not statistically significant (fig. S11A). Additionally, we measured gene expression of five interferon-related genes (*IFNA1, IFNB1, ISG15, IFI27*, and *IER3*) in SAT and VAT infected with SARS-CoV-2 over a period of 96 hpi (fig. S10). We identified increased expression of *IFNA1*, *IFNB1*, and *ISG15* as viral RNA accumulated in both SAT and VAT demonstrating induction of antiviral response (fig. S10).

To better understand the impact of SARS-CoV-2 infection on inflammatory responses within the SVC, we tested supernatants from infected versus mock-infected cultures for protein expression of various secreted inflammatory markers by Luminex. As compared to mock infection, SARS-CoV-2 infection of SAT and VAT resulted in upregulation of several chemokines, cytokines, growth factors, and other inflammatory mediators (Table 1). In both SAT and VAT, the elevated production of IP-10 is particularly noteworthy as this cytokine has been reported to be elevated in the serum of critically ill COVID-19 patients (*49*). Additionally, we found the platelet-derived growth factor (PDGF)-AA, and AB/BB (PDGFAB/BB) together with the type 2 (TH2) immune factor interleukin-4 (IL-4) to be elevated in SARS-CoV-2 infected SAT. Similarly, PDGF-AA, IL-4, and IL-13 (another TH2 factor) are elevated in the SARS-CoV-2 infected VAT. Together, these data suggest that adipose tissue could contribute to the secretion of the cytokines and vascular factors in the setting of COVID-19 infection.

**Table 1.**
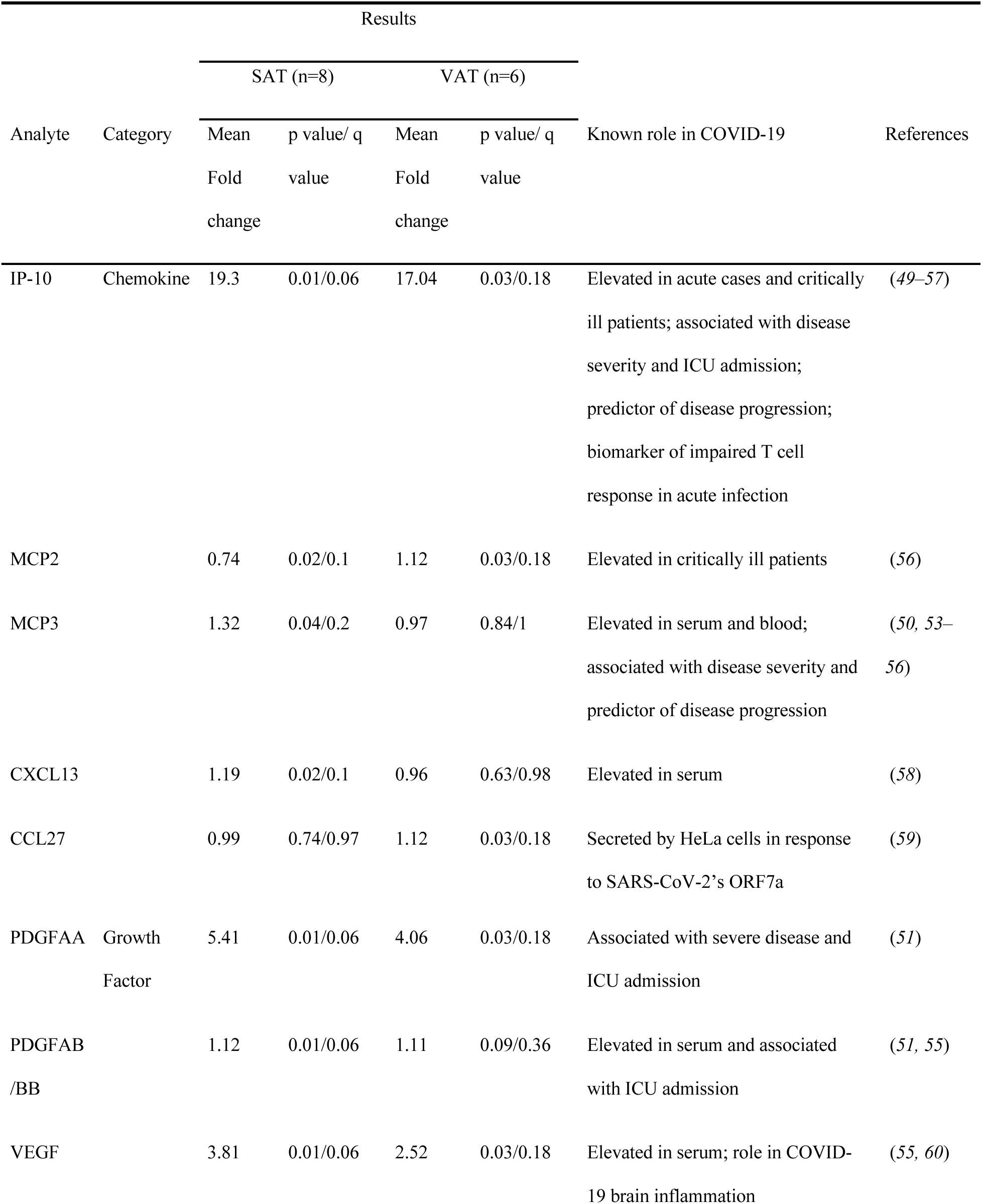
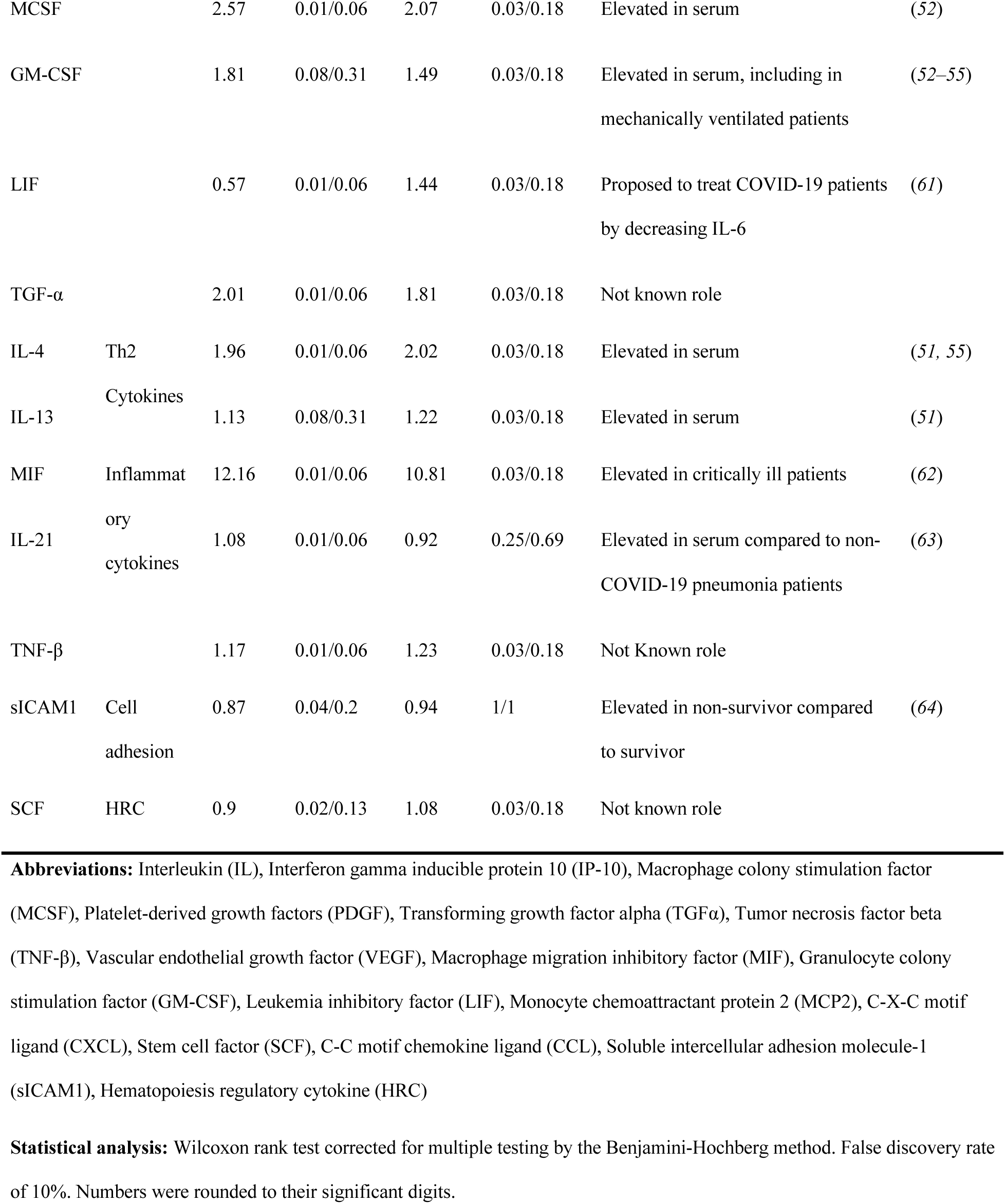
80-plex Luminex analysis of stromal vascular cells from human adipose tissue exposed to SARS-CoV-2. Table summarizing results of 80-plex Luminex assay performed in supernatants of SVC (SAT, n=8; VAT, n=6) cultures that were infected with SARS-CoV-2 or mock-infected for 24 hours. First column shows analytes of interest. Second column categorizes analytes in either chemokine, growth factor, Th2 cytokine, inflammatory cytokine, cell adhesion molecule, or hematopoiesis regulatory cytokine. Columns 3 to 6 summarizes statistical results splitted by mean fold change and p and q (Adjusted p value) value obtained from Wilcoxon signed rank test. Columns 7 and 8 summarizes the known role of each analyte in COVID-19. Bottom of the table contains abbreviations and descriptions of statistics.

### Mature and *in vitro* differentiated adipocytes support SARS-CoV-2 infection

We next explored whether mature adipocytes could support SARS-CoV-2 infection. Mature adipocytes were infected with SARS-CoV-2 or mock infected. We detected both genomic and subgenomic SARS-CoV-2 (*N gene)* in infected mature adipocytes from both SAT (n=3) and VAT (n=2 omentum; n=1 EAT; n=1 PAT) by absolute gene quantification 24 hpi (Fig. 5A). We also measured secreted inflammatory mediators by Luminex in these infected mature fat samples (fig. S9A, S9B). There was a trend for increased secretion of VEGF, PDGFAA, PDGFAB/BB and IP-10 upon infection, but no significant differences in cytokines levels (fig. S9A-B).

**Figure 5.**
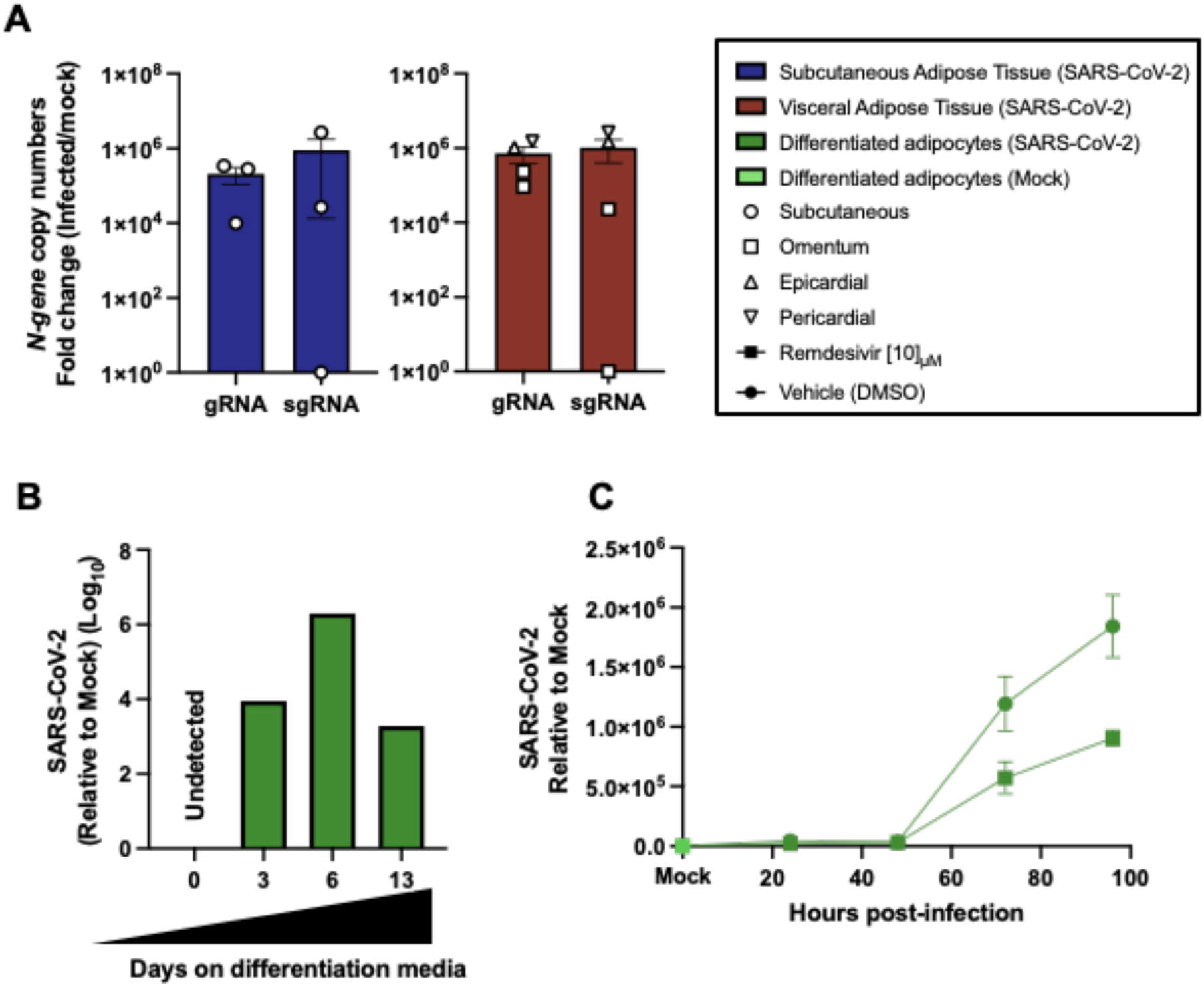
Mature and *in vitro* differentiated adipocytes support SARS-CoV-2 infection. Mature adipocytes (MA) of human adipose tissue were isolated by collagenase digestion prior to viral infection. MA were infected or left untreated (mock) for 24 hours with SARS-COV-2 (USA-WA1/2020) at a MOI:1. **(A)** Measurements of genomic (gRNA) and subgenomic (sgRNA) SARS-CoV-2 genome copy numbers infected MA from subcutaneous (left; n=3) and visceral (right; n=2 omentum; n=1 epicardial; n=1 pericardial) adipose tissue obtained by absolute gene quantification using 1-step RTqPCR TaqMan^TM^ and reported as fold change of infected to mock). **(B)** Relative gene expression of *N gene* in adipocytes differentiated *in vitro* from pericardial preadipocytes using adipocyte differentiation media for 0, 3, 6, and 13 days before left untreated or infected with SARS-CoV-2 at a MOI:1. Results were obtained by 1-step RTqPCR and analyzed by ΔΔCt method using 18s as a housekeeping gene. **(C)** Adipocytes differentiated *in vitro* from preadipocytes obtained from pericardial adipose tissue were infected or mock infected with SARS-COV-2 (USA-WA1/2020) at a MOI of 1 for 1 hour, followed by washing and removing the virus and replacing with media treated with vehicle (DMSO) or 10µM remdesivir to inhibit viral replication. Cultures were maintained for 24, 48, 72, and 96 hpi, after which gene expression was obtained by 1-step RTqPCR for the *N* gene. Relative expression was analyzed by ΔΔCt method relative to the mock sample. **(C)** Each data point is an average of 3 technical replicates and are presented as ± mean s.e.m.

We next evaluated infection in differentiated preadipocytes (*68*). As a control of successful differentiation, we confirmed an increase in gene expression of fatty acid binding protein 4 (*FABP4*) (*69*) on differentiated adipocytes (table S2B). *ACE2* expression was not detected in undifferentiated preadipocytes (day 0) but was detected by day 3 in SAT, VAT, and PAT (table S2B). Adipocytes differentiated from PAT were infected at days 3, 6, and 13 of differentiation. We detected SARS-CoV-2 (*N gene)* at 24 hpi in all differentiated adipocytes (Fig. 5B). Adipocytes differentiated from SAT had detectable genomic and subgenomic SARS-CoV-2 RNA detected 24hpi on adipocytes at day 0 and day 3 of differentiation (fig. S9C-D). To determine whether SARS-CoV-2 can replicate in differentiated adipocytes, we infected day 13 differentiated adipocytes, and cultured them with either vehicle or remdesivir. We observed increased viral reads in differentiated adipocytes by 72 hpi; this viral RNA replication was partially inhibited by remdesivir (Fig. 5C). These results show that both freshly isolated mature adipocytes and adipocytes differentiated in culture can be infected by SARS-CoV-2 and represent an additional replication site.

### SARS-CoV-2 is detected in the adipose tissue from deceased COVID-19 patients

We next sought to find evidence for SARS-CoV-2 infection of adipose tissue *in vivo*. We performed RTqPCR for SARS-CoV-2 genes in lung, fat, heart, and kidney tissue samples in a cohort of eight COVID-19 autopsy cases (Table 2; autopsy # 1-8). Studies assessing SARS-CoV-2 levels in the lungs and heart of several subjects here have been previously published (*70–72*). In addition to detecting SARS-CoV-2 in the lung and heart, we also detected SARS-CoV-2 in epicardial, visceral, and subcutaneous adipose tissue and kidney (Table 2). As expected, SARS-CoV-2 detection was highest in the lung (mean Ct=25). Interestingly, it was followed by adipose tissue (mean Ct=33), heart (mean Ct=34) and kidney (mean Ct=34). In addition, we performed RNA *in situ* hybridization (ISH) on epicardial fat (Fig. 6). We detected SARS-CoV-2 in the cytoplasm of adipocytes in the epicardium and not in immune cells or myocardium (Fig. 6A-D). Mononuclear cell infiltration, mainly consisting of lymphocytes, is observed in areas showing positive SARS-CoV-2 signal in epicardium (Fig. 6A-E). Our results provide evidence of SARS-CoV-2 detection in the adipose tissue of COVID-19 autopsies, indicating that adipose tissue tissue can harbor SARS-CoV-2 infection and contribute to pathogenic inflammation.

**Figure 6.**
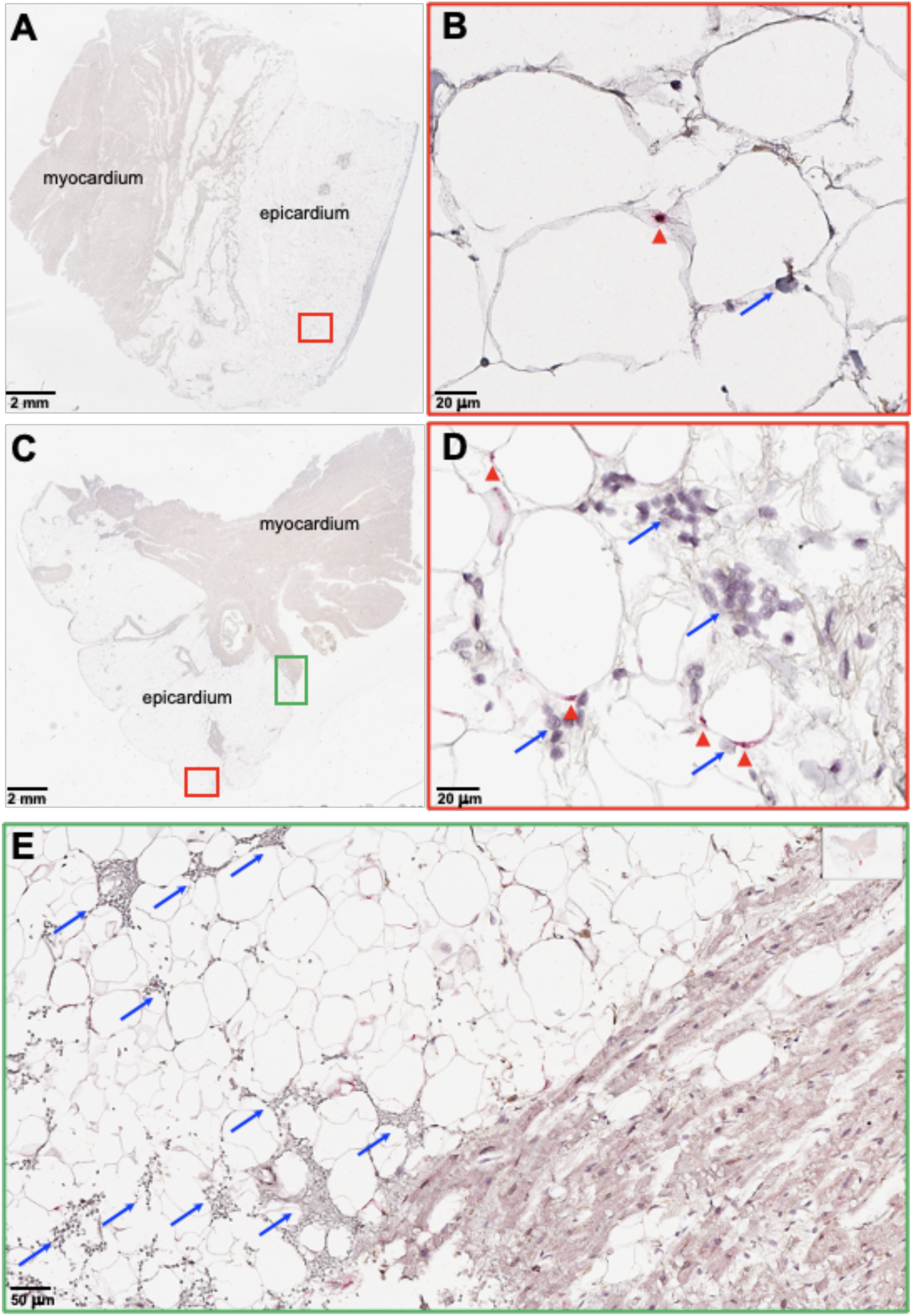
SARS-CoV-2 RNA and immune infiltration are present in adipose tissue of autopsy samples from COVID-19 patients. RNA *in situ* hybridization (ISH) on epicardial fat from heart autopsies from patients (Top, autopsy #9; Bottom, #10) who succumbed to COVID-19. Assays were performed using probes against SARS-CoV-2 Spike mRNA. Red arrowheads show ISH positive signals and blue arrows show inflammatory cells. **(A and C)** Overview of the heart tissue section (2mm), and **(B and D)** magnified view (20um) of the represented region. **(E)** Interface of epicardial fat and myocardium (50um). Note the inflammatory infiltration (blue arrows) only in the epicardial fat. Image has been rotated 90°.

**Table 2.**
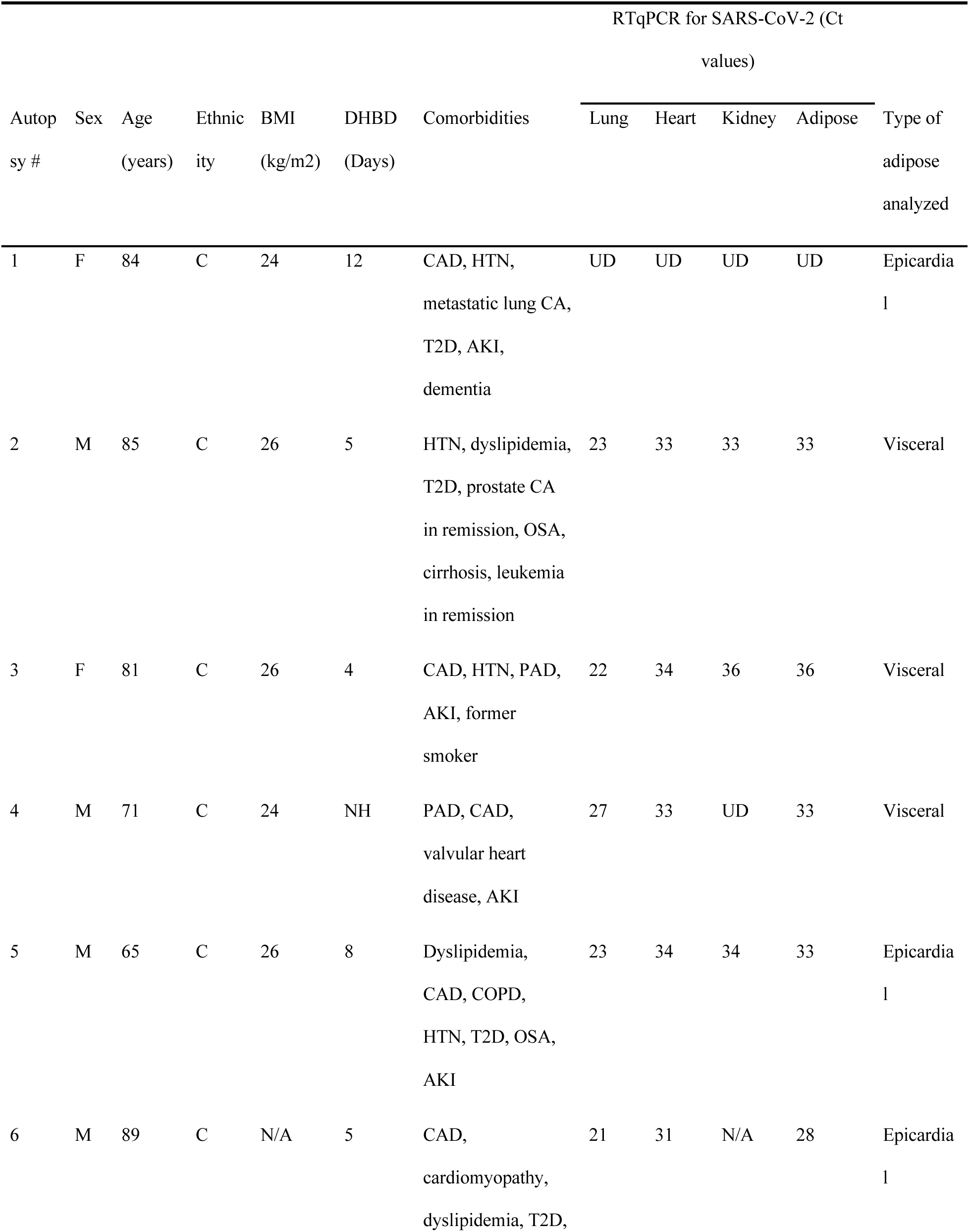
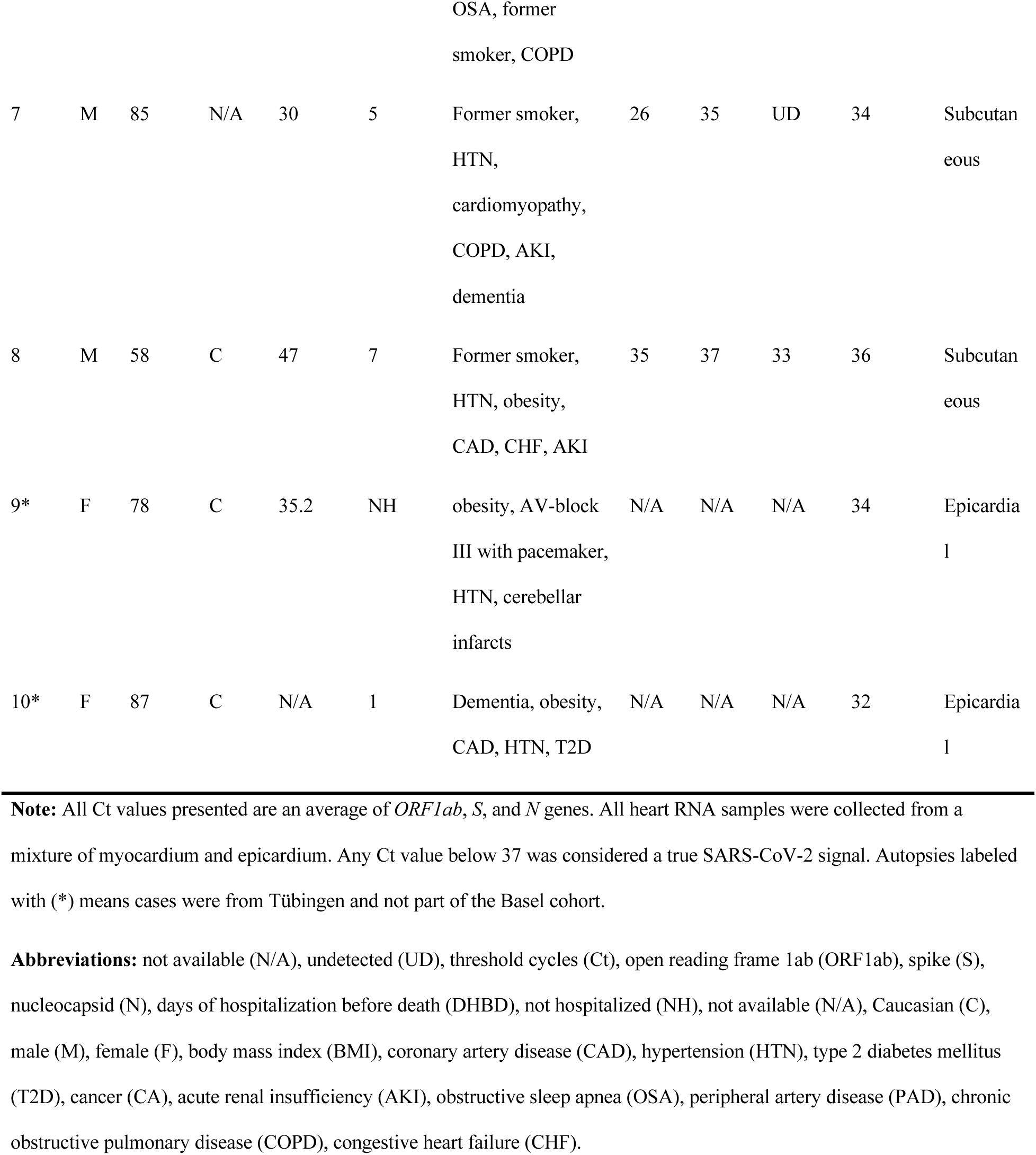
Evidence of SARS-CoV-2 detection in adipose tissue from deceased individuals. Table summarizing SARS-CoV-2 PCR signals from 8 autopsy cases, seven of them from patients deceased from COVID-19. All threshold cycles (Ct) presented are an average of three SARS-CoV-2 gene sequences for open reading frame 1ab (*ORF1ab*), spike (*S*), and nucleocapsid (*N*). RNA was isolated from either epicardial, visceral, or subcutaneous fat. Heart RNA samples were collected from a mixture of myocardium and epicardium. Samples not presented in this study were marked as not available “N/A”. When the PCR signal was undetected, the sample was marked as “UD”.

| Autopsy # | Sex | Age (years) | Ethnicity | BMI (kg/m<sup>2</sup>) | DHBD (Days) | Comorbidities | RTqPCR for SARS-CoV-2 (Ct values) |  |  |  | Type of adipose analyzed |
| --- | --- | --- | --- | --- | --- | --- | --- | --- | --- | --- | --- |
|  |  |  |  |  |  |  | Lung | Heart | Kidney | Adipose |  |
| 1 | F | 84 | C | 24 | 12 | CAD, HTN, metastatic lung CA, T2D, AKI, dementia | UD | UD | UD | UD | Epicardial |
| 2 | M | 85 | C | 26 | 5 | HTN, dyslipidemia, T2D, prostate CA in remission, OSA, cirrhosis, leukemia in remission | 23 | 33 | 33 | 33 | Visceral |
| 3 | F | 81 | C | 26 | 4 | CAD, HTN, PAD, AKI, former smoker | 22 | 34 | 36 | 36 | Visceral |
| 4 | M | 71 | C | 24 | NH | PAD, CAD, valvular heart disease, AKI | 27 | 33 | UD | 33 | Visceral |
| 5 | M | 65 | C | 26 | 8 | Dyslipidemia, CAD, COPD, HTN, T2D, OSA, AKI | 23 | 34 | 34 | 33 | Epicardial |
| 6 | M | 89 | C | N/A | 5 | CAD, cardiomyopathy, dyslipidemia, T2D, | 21 | 31 | N/A | 28 | Epicardial |

Table 2. Evidence of SARS-CoV-2 detection in adipose tissue from deceased individuals.
|  |  |  |  |  |  |  |  |  |  |  |  |
| --- | --- | --- | --- | --- | --- | --- | --- | --- | --- | --- | --- |
|  |  |  |  |  |  | OSA, former<br>smoker, COPD |  |  |  |  |  |
| 7 | M | 85 | N/A | 30 | 5 | Former smoker,<br>HTN,<br>cardiomyopathy,<br>COPD, AKI,<br>dementia | 26 | 35 | UD | 34 | Subcutan<br>eous |
| 8 | M | 58 | C | 47 | 7 | Former smoker,<br>HTN, obesity,<br>CAD, CHF, AKI | 35 | 37 | 33 | 36 | Subcutan<br>eous |
| 9* | F | 78 | C | 35.2 | NH | obesity, AV-block<br>III with pacemaker,<br>HTN, cerebellar<br>infarcts | N/A | N/A | N/A | 34 | Epicardia<br>l |
| 10* | F | 87 | C | N/A | 1 | Dementia, obesity,<br>CAD, HTN, T2D | N/A | N/A | N/A | 32 | Epicardia<br>l |

**Note:** All Ct values presented are an average of *ORF1ab*, *S*, and *N* genes. All heart RNA samples were collected from a mixture of myocardium and epicardium. Any Ct value below 37 was considered a true SARS-CoV-2 signal. Autopsies labeled with (\*) means cases were from Tübingen and not part of the Basel cohort.
**Abbreviations:** not available (N/A), undetected (UD), threshold cycles (Ct), open reading frame 1ab (ORF1ab), spike (S), nucleocapsid (N), days of hospitalization before death (DHBD), not hospitalized (NH), not available (N/A), Caucasian (C), male (M), female (F), body mass index (BMI), coronary artery disease (CAD), hypertension (HTN), type 2 diabetes mellitus (T2D), cancer (CA), acute renal insufficiency (AKI), obstructive sleep apnea (OSA), peripheral artery disease (PAD), chronic obstructive pulmonary disease (COPD), congestive heart failure (CHF).

## DISCUSSION

Here, we report that SARS-CoV-2 infects human adipose tissue in COVID-19 patients and *in vitro.* Significantly extending a prior report that human adipocytes can be infected *in vitro* (*15*), we identify both tissue-resident macrophages and mature adipocytes as target cells for SARS-CoV-2 infection (*16, 17, 19*). Additionally, this study reveals that *in vitro* infection leads to activation of inflammatory pathways in macrophage and preadipocytes and the secretion of inflammatory factors associated with severe COVID-19. Finally, we provide the first evidence that this may be relevant to disease pathogenesis in humans *in vivo*, as we detected SARS-CoV-2 RNA in VAT, EAT, and SAT of COVID-19 autopsies. We also demonstrate histologic evidence of inflammation adjacent to viral signals in adipose tissue in an autopsy sample. Together, this in-depth analysis of adipose susceptibility and inflammatory response to SARS-CoV-2 infection suggests that adipose tissue may serve as a potential reservoir for SARS-CoV-2 and potentiator of systemic and regional inflammation, possibly contributing to severe clinical disease in obese individuals infected with SARS-CoV-2.

One of our most intriguing findings was that SARS-CoV-2 infection of adipose SVC was primarily restricted to just one of two macrophage clusters. The two macrophage populations were defined as: C2-macrophages, an inflammatory cluster with *IL1B* and *CCL3* as its most distinguishing transcripts, and C12-macrophages, characterized by significant enrichment of HLA class II transcripts and *LYZ* and alarmin *S100A8* expression. C2-macrophages were predominately infected, with over 85% of C2 cells within infected SVC carrying SARS-CoV-2 transcripts. These two clusters do not fall into a classical M1 and M2 classification: instead, macrophages activated in response to adipose tissue signals may exhibit a unique phenotype (*39, 73*). The C2 macrophages express transcripts characteristic of PVMs, which have been previously described in adipose tissue (*31*) and are highly phagocytic (*74*). PVMs can become virally infected: in the brain, PVMs have been shown to become infected by HIV and SIV (*38, 75*). Interestingly, SARS-CoV-2 infection further drives the C2/C12 dichotomy as chemokine markers are further upregulated while *LYZ* and HLA expression is further downregulated in infected SARS-CoV-2 transcript-containing C2 cells versus their bystander counterparts. *LYZ* is of particular interest as prior studies have shown that lysozyme can play a role in antiviral activity (*76–78*), which suggests that low lysozyme levels (as found in the susceptible C2 cluster) may be advantageous for the virus. For example, influenza infection reduces lysozyme secretion in neutrophils (*79*) and the combination of lysozyme and lactoferrin therapy reduces Bovine viral diarrhoea virus titers *in vitro* (*76, 78*). This data suggests that lysozyme therapy could reduce SARS-CoV-2 levels in the adipose tissue. We also observed significant reduction in HLA Class I and II genes relevant to antigen presentation in the infected C2-macrophages cluster, consistent with findings in blood monocytes in severe COVID-19 disease (*80, 81*) which reduces immune surveillance and targeting of infected host cells, another infection-induced change which could be advantageous for virus propagation. Together, these data highlight the pivotal role macrophages may play in disease pathogenesis and the need for a better understanding of how tissue macrophages at different sites are infected by and control SARS-CoV-2 infection.

The role of macrophages in supporting SARS-CoV-2 replication has been a subject of significant interest as there is no consensus about whether macrophages support viral replication. SARS-CoV-2 and the related coronaviruses SARS-CoV-1 and MERS can enter macrophages (*82–84*), but the downstream effects remain unclear. One study shows that neither MERS-CoV nor SARS-CoV-1 can replicate in human macrophages *in vitro* (*82*), another found that MERS-CoV but not SARS-CoV-1 can replicate within human monocyte-derived macrophages (*83*), and yet another study demonstrated abortive SARS-CoV-1 infection of human macrophages (*84*). For SARS-CoV-2, a recent study reported that the virus fails to replicate or induce an inflammatory response in human monocyte-derived macrophages *in vitro* (*85*), while another study showed that the SARS-CoV-2 spike protein can induce macrophage activation in murine cells (*86*). Importantly, these studies focused on blood-derived macrophages, and tissue-resident macrophages may have different properties. SARS-CoV-2 has been detected in human alveolar macrophages COVID-19 patients (*87*). While it is possible that infection in adipose tissue macrophages is abortive, two lines of evidence suggest that this infection is productive. First, we detected both genomic RNA and positive sense sgRNA, an intermediate RNA made during viral replication (*88*); however, these data should be interpreted with caution as sgRNAs can be quite stable and therefore may not fully reflect active replication (*89, 90*). Second, we observed increased viral RNA accumulation over time in SVC infections, suggestive of viral replication, which was decreased by remdesivir, an inhibitor of the viral RNA-dependent RNA polymerase. Interestingly, we also observed enrichment in inflammatory pathways and a reduction in host translation pathways specifically in infected macrophages, indicative of viral hijacking of host translation (*91*). Such suppression of host translation has been also noted in SARS-CoV-1 infection: the NSP1 protein of SARS-CoV-1 suppresses host translation and gene expression, including those of type I interferons (*43*). Together, these data highlight the dramatic effects of SARS-CoV-2 infection on macrophages.

We also found that SARS-CoV-2 can infect human adipocytes both *in vitro* and *in vivo*. Previously, researchers demonstrated that SARS-CoV-2 is detectable in adipose tissue of SARS-CoV-2-infected hamsters, and that human adipocytes derived from breast tissue can support infection *in vitro* (*15*). Here we demonstrate infection in multiple different adipose depots, including critical peri-organ depots. We show this through (1) detection of both genomic and subgenomic SARS-CoV-2 *N gene* expression in infected mature adipocytes and (2) infected differentiated preadipocytes, (3) demonstration of a reduction in SARS-CoV-2 viral load in differentiated preadipocytes post-infection in response to remdesivir treatment, and (4) histology showing SARS-CoV-2 in adipocytes in a COVID-19 autopsy. This data adds to a growing body of evidence that adipose tissue can serve as a reservoir for RNA viruses including influenza A virus and HIV (*16, 17, 19*). Collectively, this data suggests that adipose tissue may serve as a reservoir for SARS-CoV-2.

Our data show that preadipocytes adopted proinflammatory phenotypes following SARS-CoV-2 infection of SVC, despite not being infected themselves. We noted 17 different preadipocyte clusters that varied in their representation among VAT and SAT depots. Interestingly, we found that adipocyte stem cell-like clusters demonstrated more subdued transcriptional changes in response to infection compared to other clusters across depots and participants. Pathways upregulated across various preadipocyte clusters in response to infection included interferon and interleukin signaling, both of which are important to controlling viral infection. Previous reports have demonstrated that IL-6 and TNF-ɑ cytokine exposure can impair preadipocyte differentiation and lipid accumulation, instead promoting the transition of preadipocytes towards an inflammatory state (*44, 92*). Thus, the SARS-CoV-2 infection of adipose tissue could drive a proinflammatory cascade promoted by preadipocyte activation. Interestingly, SAT preadipocytes had a more dramatic inflammatory response than VAT preadipocytes. This was surprising because several studies have pointed to the VAT being a better predictor of COVID-19 severity than the SAT (*93–95*). For example, a recent single-center cohort study demonstrated that increased VAT thickness and lower SAT thickness increased the risk of COVID-19 ICU admission, independent of BMI (*95*). While bulk transcriptomic data collected from *in vivo* SARS-CoV-2 infection of hamsters concluded that VAT had a stronger antiviral response than SAT, this study included only mature adipocytes and studied the response 48 hpi (*15*). While our human data similarly demonstrates higher cytokine expression of VAT preadipocytes at baseline, it also suggests that the SAT preadipocytes might play an underappreciated role early upon infection by driving an increased inflammatory response.

A significant finding of our study is the dramatic inflammatory response following SARS-CoV-2 infection of adipose tissue, particularly within the SVC. Across multiple experiments, at both the RNA and protein level, we saw increased expression of cytokines, ISGs and other molecules related to inflammation and antiviral pathways 24 hours after SVC infection. Many of the molecules we found upregulated upon SARS-CoV-2 infection are associated with COVID-19 severity: IP-10, PDGFAA, PDGFAB/BB, IL-4, and MCSF all have been reported to be elevated in the serum of critically ill COVID-19 patients (*49, 51, 52*), MIF has been described as a predictor of poor outcome on mechanically ventilated COVID-19 patients (*62*), and VEGF may play a crucial role in COVID-19 related brain inflammation (*60*). Further, we detected increased type I interferon transcripts and *ISG15* in SVC of both SAT and VAT 72 and 96 hpi, indicative of a persistent antiviral response during viral replication. While significantly less dramatic, we also saw trends for increased inflammatory cytokine and chemokines following infection of mature adipocytes. Notably, we observed both induction of transcripts (CCL8 and CCL3) and demonstrated secretion of chemokines (MCP2 and MCP3) in infected SVC cultures that can attract macrophages. This is entirely consistent with our autopsy finding of an inflammatory infiltrate associated with SARS-CoV-2^+^ cells and suggests that the antiviral response may be dragging more susceptible macrophages to the site of infection. These data suggest that targeted inhibition of inflammation in the adipose tissue could improve outcomes in COVID-19 subjects. In fact, therapeutics targeting inflammation of the adipose tissue during obesity-induced inflammation have been proposed as a treatment for metabolic disease (*96*). For instance, salicylate, a cyclooxygenase inhibitor, reduces inflammation of the adipose tissue in obese individuals and has been proposed as a therapeutic strategy in COVID-19 patients due to its anti-inflammatory properties and antiviral activity against both DNA and RNA viruses (*97–103*). Therefore, drugs reducing inflammation of the adipose tissue in obese individuals should be further explored in COVID-19 subjects.

The mechanism(s) of viral entry into adipocytes and SVC macrophages remain unclear but may not be through the canonical ACE2 receptor. While *ACE2* mRNA expression has been reported in adipose tissue, (*113*) and its level is affected by diet and obesity (*114*), we detected no ACE2 protein in SVC from VAT, and almost no expression in SAT. We could not detect *ACE2* gene expression in SVC from either SAT or VAT, either by RTqPCR with primers validated on kidney tissue or by scRNA-seq. Similarly, we did not detect *ACE2* in mature adipocytes, though we did identify upregulation of *ACE2* transcripts during *in vitro* differentiation of preadipocytes. Another recent study also detected higher levels of *NRP1* and *FURIN* (two other proposed SARS-CoV-2 viral entry factors) compared to *ACE2* and *TMPRSS2* in murine and human adipocytes (*15*). In both macrophages and adipocytes, entry could occur via use of an alternate entry receptor. Genome-scale CRISPR loss-of-function screens in susceptible human cells have identified multiple SARS-CoV-2 entry factors in addition to *ACE2* (*115*). Alternative entry receptors reported include CD147, Neuropilin-1, Dipeptidyl peptidase 4 (DPP4), alanyl aminopeptidase (ANPEP), glutamyl aminopeptidase (ENPEP), and angiotensin II receptor type 2 (AGTR2) (*104, 105*). In macrophages, entry could also occur via phagocytosis of viral particles, phagocytosis of infected cells, or, if antibodies were present, via antibody-dependent entry. Elucidating the entry mechanisms will be an important area of future investigation given its implications for disease pathogenesis. For example, Neuropilin-1 has may mediate entry into neural cells and play a role in neurologic manifestations of COVID-19 (*106*), and yet antibody therapies that only block interactions with ACE2 may not block such entry if it is relevant *in vivo*.

Our study has several limitations. Our numbers of replicates were limited for some assays, such as the evaluation of cytokine secretion following infection of mature adipocytes. Nonetheless, we observed significant indications of inflammatory responses. It is possible that we were unable to fully wash input virus following infection of mature adipocytes due to the high lipid content, size, and fragility of the cells, falsely increasing the viral signal. However, our detection of sgRNA, the time-dependent increase in viral RNA accumulation that was inhibited by remdesivir, and the detection of SARS-CoV-2 RNA in autopsy samples all provide orthogonal support for true infection of adipocytes. Our autopsy studies were limited in number, and we were only able to perform confirmatory ISH on epicardium, not in the subcutaneous, omental, or pericardial fat due to limited autopsy tissue availability. All experiments were performed with the WA-01 strain of SARS-CoV-2 and no experiments with its variants were performed, and plaque assays to confirm viral production were not performed. As tissue donors were obese, an important area of future investigation will be to study the effects of SARS-CoV-2 infection on lean adipose tissue, as well as to study the adipose tissue of those with ‘long COVID’.

Overall, here we provide evidence that two cell types within human adipose tissue are permissive to SARS-CoV-2 infection. This adds to data showing susceptibility of other tissues including heart, kidneys, pharynx, liver, brain, and pancreas (*1, 27, 107*). SARS-CoV-2 RNA was detected in autopsy samples at a higher viral load than in adjacent organs, the kidney or heart. Importantly, infection of adipose tissue drives an inflammatory response in infected macrophages and preadipocytes. Thus, multiple cells within adipose tissue likely participate in both viral replication and inflammation. Importantly, we demonstrated infection and inflammation in adipose tissue adjacent to critical organs such as the heart and intestine, thus pointing to the potential for adipose tissue potentiation of organ damage in severe COVID-19. Furthermore, if adipose cells constitute a reservoir for viral infection, obesity may contribute not only to severe acute disease, but also to long-COVID syndrome. Collectively, our data implies that infection in adipose tissue may partially explain the link between obesity and severe COVID-19. More efforts to understand the complexity and contributions of this tissue to COVID-19 pathogenesis are warranted.

## MATERIALS AND METHODS

### Study design

The aim of this study was to determine if human adipose tissue is permissive to SARS-CoV-2 infection. We obtained adipose tissue from consented subjects and exposed these samples to SARS-CoV-2 and measured viral entry, replication, and inflammatory pathways by flow cytometry, RTqPCR, scRNA-seq, and Luminex. All analyses were performed in an unbiased fashion. Formalin-fixed and paraffin embedded lung, kidney, adipose and heart tissue from COVID-19 autopsy samples were evaluated for viral RNA. The n value was not controlled and was dependent on availability of samples.

### Subjects

Study participants were recruited from the Adult Bariatric Surgery and Cardiothoracic Surgery clinics at Stanford University Medical Center during the preoperative visit. Eligibility requirements include >25 yrs of age, and exclusion criteria included chronic inflammatory conditions, pregnancy/lactation, use of weight loss medications, and current/prior diagnosis of COVID-19. The protocol was approved by the Stanford Institutional Review Board and all subjects gave written, informed consent. Tissue samples from COVID-19 deceased patients were obtained from University Hospital Tübingen, Tübingen, Germany, or from University Hospital Basel or Cantonal Hospital Baselland, Switzerland. SARS-CoV-2 infected tissue was obtained during autopsy and processed as previously described (71). The use of SARS-CoV-2 infected tissue was approved by the ethics commission of Northern Switzerland (EKNZ; study ID: 2020-00969). All COVID-19 patients or their relatives consented to the use of tissue for research purposes.

### Preparation, isolation, and differentiation of adipose tissue

On the day of bariatric surgery, approximately 2-3g each of subcutaneous abdominal (SAT) and omental visceral adipose (VAT) tissue was harvested intraoperatively and immediately processed. For cardiothoracic surgery patients, 2g of PAT, 1g of EAT, and 1-2g of SAT chest wall was obtained intraoperatively and immediately processed. Tissue was subject to collagenase digestion for separation of mature adipocytes (MA) and SVC, and for differentiation of preadipocytes as previously described (108) with details in supplemental methods.

### Virus and cell lines

The USA WA1/2020 strain of SARS-CoV-2 was obtained from BEI Resources, passaged in VeroE6 cells, and tittered by Avicel (FMC Biopolymer) overlay plaque assay on VeroE6 cells. Passage 3 virus was used for all infections. VeroE6 cells were obtained from ATCC and were mycoplasma free. A549-ACE2 was a gift from Ralf Bartenschlager and was mycoplasma free.

### SARS-CoV-2 infections of differentiated adipocytes, SVC, MA, and A549-ACE2

Cells were infected with SARS-CoV-2 (2019-nCOV/USA-WA1/2020) at a MOI of 1 for 1 hour before washing input virus and replacing media for incubation. Remdesivir was used at 10 µM.

### RNA isolation and quantification

At the time of collection, cells were washed with PBS followed by incubation with TRIzol LS (Thermo) reagent for 15-20mins for cell lysis and virus inactivation and RNA extraction. RNA was resuspended in water and quantified by absorbance (260/280) using a NanoDrop^TM^ spectrophotometer (Thermo Scientific).

### Genomic and subgenomic absolute gene quantification by RTqPCR

5µl of the total RNA was used for 1-step RTqPCR. Genomic N-gene quantification was done with the use of CDC qualified primers and probes amplifying the N1 region, n2019-nCoV (Biosearch technologies, KIT-NCOV-PP1-1000). For subgenomic *N-gene* quantification, *E-gene* sgRNA forward primer for SARS-CoV-2 leader sequence was combined with CDC N1 gene reverse primer and probe to detect N-gene sgRNA as previously shown (*89, 109*). RNA and primers were mixed with the TaqPath 1-step RTqPCR master mix (Applied Biosystems, A15299). A standard curve for Ct values and genome copy numbers was obtained using pET21b+ plasmid with the N-gene inserts. All the samples were analyzed in technical duplicates. The Ct cutoff for positive samples was <38 with amplification observed in both duplicates. The samples were analyzed on a StepOnePlus^TM^ real time PCR machine (Applied Biosystems) using the following parameters: (stage 1) 10 minutes at 50°C for reverse transcription (RT), followed by (stage 2) 3 minutes at 95°C for initial denaturation and (stage 3) 40 cycles of 10 seconds at 95°C, 15 seconds at 56°C, and 5 seconds at 72°C.

### RTqPCR for relative gene quantification

RNA was isolated as described above. Either the 1- step method using TaqPath (Applied Biosystems, A15299) or the 2-step process using the Superscript III first-strand synthesis system was performed (Applied biosystems, A25742) (for details see the Supplemental Materials and Material).

### SARS-CoV-2 detection in autopsies

Autopsy samples were prepared as previously described (110). Briefly, RNA was isolated from formalin-fixed and paraffin embedded tissue with the use of Maxwell RSC RNA FFPE (Promega) according to manufacturer recommendations. RTqPCR was performed using the TaqMan^TM^ 2019-nCoV control kit v1 (Thermo Scientific, A47533) to target the three viral genes: *ORF1ab*, *S*, and *N* genes, and the human *RPPH1* gene. According to the manufacturer’s recommendation, a Ct value below 37 in at least two out of three viral genomic regions was considered positive. A case was considered negative if Ct values were above 40. Values between 37 and 40 were considered undetermined and the assay was repeated. Samples were always run in duplicates. RNA-ISH was used to detect SARS-CoV-2 Spike mRNA in tissue samples (for details see the Supplemental Materials and Material).

### Flow cytometry

A single cell suspension was stained with Fc block, a viability stain, and surface stained for CD45, CD3, CD14, CD34, CD11c, and CD31 before fixing and permeabilizing to stain with anti-SARS-CoV-2 N protein as described in supplemental methods.

### Sample preparation for Luminex

Supernatants from mock or infected SVC were collected after 24 hpi into low-binding protein collection tubes. Supernatants from mature adipocytes were collected by first removing floating mature adipocytes with a cut wide pipette tip and pipetting remaining media into low-protein binding collection tubes. Supernatants were stored at -80°C. To remove samples from BSL3 containment, supernatants were thawed and mixed with 10% TritonX-100 (Sigma-Aldrich, T9284) for a final concentration of 1% TritonX-100 and incubated for 20 minutes at room temperature for viral inactivation. Supernatants were then removed from BSL3 and transferred to a low-binding protein 96 well plate for 80plex Luminex by the Human Immune Monitoring Center (HIMC) at Stanford University.

### Single cell RNA sequencing (scRNA-seq)

The gel beads-in-emulsion (GEM) single cell 3’ kit v3.1 dual index kit (10X Genomics) was used following manufacturer’s recommendations with slight modifications. Briefly, SVC from SAT and VAT was left untreated or infected with SARS-CoV-2 as described above. After 24hpi, cells were collected, washed, and diluted at a density of 1×10^3^ cells per µl in cold DMEM (Life technologies; 11885-092) media supplemented with 10% FBS (Corning, MT35016CV). 10,000 cells per lane were loaded onto a Chromium Controller in the BSC per manufacturer’s instructions. Following GEM creation, samples were transferred into PCR tube strips prior to transferring into a PTC-200 thermocycler (MJ Research) for RT. The RT parameters were the following: 45 minutes at 53°C, followed by 5 minutes at 85°C, then 15 minutes at 60°C and finally samples were kept at 4°C. Barcoded cDNA was removed to BSL2, and sequencing libraries were prepared per manufacturer’s recommendation, with a TapeStation 4200 (Agilent) used for quality control. Libraries were pooled for a final concentration of 10nM and sequenced on a NovaSeq S2 (Ilumina) at the Chan Zuckerberg Biohub (San Francisco).

### Alignment and preprocessing of scRNA-seq data

The quality of the raw FASTQ data was examined with FASTQC and then aligned (cell ranger count) to a custom genome including human genome (hg38) and the complete genome sequence of SARS-CoV-2 (2019-nCOV/USA-WA1/2020) (GenBank: MN985325.1) using the “Cell Ranger” software package v6.0.0 (10x Genomics). R (4.0.4) was used for all downstream analyses. Resulting filtered feature-barcode-matrices were processed using the R package Seurat (v4.0.0). Briefly, count matrices were merged and loaded into Seurat with SARS-CoV-2 counts removed and appended to the metadata. All genes represented in < 10 cells were excluded from downstream analysis. Cells were filtered based on the following criteria: less than 200 distinct genes, less than 100 unique molecular identifiers (UMIs), and greater than 15% UMIs from mitochondrial genes. Each batch was then individually normalized using the “SCTransform” function that included regression for percent mitochondria. Integration features were then calculated using the “SelectIntegrationFeatures” function and passed into “VariableFeatures” of the merged object to maintain the repeatedly variable features across each dataset. Within each batch and condition (tissue and infection status), cells identified as doublets by both “DoubletFinder” (v3) (using pN = 0.25, pK = 0.09, PCs = 1:50, anticipated collision rate = 10%) and scds (top 10% of cells ranked by hybrid scores) were removed (n = 3,165 cells) from the analysis. After applying these filtering steps, the dataset contained 198,759 high-quality cells. Principal component (PC) analysis (PCA) was performed on the data. The resulting data were corrected for batch effects using the “Harmony” package (*29*) with the top 50 PCs. Uniform Manifold Approximation and Projection for Dimension Reduction (UMAP) coordinate generation and clustering were performed using the “RunUMAP”, “FindNeighbors”, and “FindClusters” (resolution = 0.6) functions in Seurat with PCs 1-50. Manual annotation of each cellular cluster was performed by finding the differentially expressed genes using Seurat’s “FindAllMarkers” function with default Wilcoxon rank-sum test and comparing those markers with known cell type-specific genes from previous datasets (*30, 31*). “FindMarkers” function using the MAST algorithm (latent.vars = ‘participant’) based on a Bonferroni-adjusted P < 0.05 and a log2 fold change > 0.25 was used for targeted differential gene expression analysis.

### scRNA-seq analyses

Gene ontology and KEGG pathway analysis was performed using R package stringdb (*111*). Reactome gene set enrichment analysis was performed using the R package fgsea (*112*), considering ranked gene lists. The Seurat function AddModuleScore() was used to score individual cells by expression of either a list of genes relating to ISGs or cytokines. This function generates an average module score by calculating the mean expression of each gene in the module corrected for expression of a random set of similarly expressing genes. Gene lists used to define each module are defined in table S15. Heatmaps were generated using ComplexHeatmap, Seurat and pheatmap packages, Violin plots were generated using ggplot2, and dotplots were generated using the Seurat packages in R. A Github repository for all original code used for analysis and visualization will be made public upon publication.

### Images

Pictures were taken with the use of a EVOS XL core cell imaging system (Thermo Fisher Scientific) with an objective of 10x.

### Statistical analysis

GraphPad Prism version 9.1.0 (216) and R (4.0.4) were used for statistical analysis. When comparing mock and infected groups a paired, two-sided, student’s t-test was applied. When comparing more than two groups a two-way ANOVA, multiple comparisons using statistical hypothesis Sidak was performed. In Luminex analysis a paired Wilcoxon signed rank test with desired false discovery rate of 10% was performed. Data bars are always presented as ± mean s.e.m.

## Supplementary Materials

### Materials and Methods

Fig. S1. Genomic and Subgenomic measurement of SARS-CoV-2 on in vitro infected SVC.

Fig. S2. Validation of Remdesivir treatments in A549-ACE2 cells.

Fig. S3. Gating strategy of SVC in human adipose tissue.

Fig. S4. Limited ACE2 protein detection in SVC from SAT and VAT.

Fig. S5. Infection, depot and participant breakdown by cluster annotation.

Fig. S6. Distribution of SARS-CoV-2 transcripts across all cells.

Fig. S7. C2- and C12-macrophages are distinctly different upon mock and SARS-CoV-2 infection.

Fig. S8. Characterization of preadipocytes across VAT and SAT.

Fig. S9. Mature and *in vitro* differentiated adipocytes harbor SARS-CoV-2 RNA and mount inflammatory responses after exposure to SARS-CoV-2 *in vitro*.

Fig. S10. Increased interferon related genes in adipose tissue post-SARS-CoV-2 infection.

Fig. S11. Elevated IL-6 gene expression and secretion of inflammatory mediators in SVC post in vitro SARS-CoV-2 infection.

Table S1. Adipose tissue participant’s demographic, medical information, and sample use.

Table S2. No expression of ACE2 in adipose tissue but increased ACE2 expression in in vitro differentiated adipocytes at 3 days of differentiation.

Table S3. Marker genes for each cluster defined in combined Seurat analysis.

Table S4. Differentially expressed transcripts within macrophages only, across different subsets of infection, mock and depot.

Table S5. Differentially expressed transcripts between infected (SARS-CoV-2+) versus bystander populations within the C2 and C12 infected macrophage populations.

Table S6. GO and KEGG term enrichment for markers of C12 macrophages.

Table S7. GO and KEGG term enrichment for markers of C2 macrophages.

Table S8. Reactome pathways for differentially expressed genes between infected (SARS-CoV-2+) versus bystander C2 macrophages.

Table S9. Marker genes for each cluster defined from the Seurat analysis of Preadipocytes only.

Table S10. Differentially expressed genes between SARS-CoV-2 exposed versus mock exposed preadipocytes, by cluster and participant, within the SAT

Table S11. Differentially expressed genes between SARS-CoV-2 exposed versus mock exposed preadipocytes, by cluster and participant, within the VAT.

Table S12. Reactome pathways for differentially expressed genes between SARS-CoV-2 exposed versus mock exposed preadipocytes, by cluster and participant, within the SAT.

Table S13. Reactome pathways for differentially expressed genes between SARS-CoV-2 exposed versus mock exposed preadipocytes, by cluster and participant, within the VAT.

Table S14. ISG and cytokine gene modules.

## Acknowledgements

We thank the patients and their families for their consent to use their tissues. We would like to thank the Stanford Bariatric Surgery and Cardiothoracic Surgery clinic staff for assisting with participant recruitment and tissue harvesting. We thank Dr. Karin Klingel, Dr. Selina Traxler, Karen Greif, Dr. Massimo Granai and Dr. Hans Bösmüller (Department of Pathology, University Hospital Tübingen, Germany) for providing samples and logistic support. We would like to thank Anna Stalder and Jan Schneeberger (Institute of Medical Genetics and Pathology, University Hospital of Basel, Switzerland) for data analysis and technical assistance. We are thankful to Ralf Bartenschlager (University of Heidelberg, Germany) for providing A549-ACE2. The following reagent was deposited by the Centers for Disease Control and Prevention and obtained through BEI Resources, NIAID, NIH: SARS-Related Coronavirus 2, Isolate USA-WA1/2020, NR-52281. We thank Yael Rosenberg-Hasson, technical director of immunological assays at Stanford University’s HIMC, for Luminex assays. We thank Jaishree Garhyan, director of Stanford’s BSL3 service center, for BSL3 surveillance and training. We are grateful to Angela Detweiler and the Chan Zuckerberg Biohub foundation for sequencing.

## Funding

National Institutes of Health grant R21AI159024 (TLM)

American Diabetes Association grant 7-20-COVID-213 (TLM)

Stanford University Innovative Medicines Accelerator COVID-19 Response grant (CAB, TLM)

Botnar Research Centre for Child Health grant Emergency Response to COVID-19 grant (SJ, CMS, KDM, AT, MSM, GPN)

Swiss National Science Foundation grant 320030_189275 (MSM)

Chan Zuckerberg Biohub Investigator Program (CAB)

National Institutes of Health grant 5T32 AI007502 (AR)

National Science Foundation Graduate Research Fellowship 2019282939 (KR)

Bill and Melinda Gates Foundation OPP1113682 (JRA)

## Author contributions

Conceptualization: GMC, HeC, CAB, TLM

Methodology: GMC, KR, HeC, SJ, HaC, CAB, TLM

Investigation: GMC, KR, HeC, SZ, SJ, CMS, MSM, HaC, RV, AR

Resources: EZ, DA, JB, AT, KDM, MSM, CMS, CAB, TLM

Data Curation: GMC, KR, CAB, TLM

Visualization: GMC, KR, SZ, SJ, CMS, MSM

Funding acquisition: CAB, TLM, SJ, CMS, AT, KDM, MSM, GPN

Supervision: JRA, GPN, CAB, TLM

Writing – original draft: GMC, KR, HeC, CAB, TLM

Writing – review & editing: GMC, KR, HeC, SJ, EZ, AR, RV, HaC, JRA, DA, JB, GPN, CMS, MSM, CAB, TLM

## Data and materials availability

Data from scRNA-seq will be deposited with the Gene Expression Omnibus. A Github repository for all original code used for analysis and visualization will be made public upon publication.

## SUPPLEMENTARY MATERIAL

### Materials and Methods

#### Adipose Tissue Processing

Adipose tissue samples were minced and then digested by collagenase I (1 mg/mL) (Worthington Biochem. Corp, USA) at 37**°**C for 60 minutes, in KRBH buffer containing BSA (2%), adenosine (250 uM), and P/S (1x), then filtered through a 500-µm nylon mesh, followed by separation of MA and SVC. MA were utilized immediately; SVC was collected by centrifugation at 500 xg for 5 minutes at room temperature. The SVC pellet was incubated in erythrocyte lysis buffer (Invitrogen) for 10 minutes at 37**°**C, followed by centrifugation as above. The cell pellet was resuspended in HBSS (with 2% BSA) and centrifuged for another 5 minutes at 500 xg at RT. The SVC pellets were resuspended in growth medium (DMEM/F12, 10% FBS, and 1% P/S), filtered through a 75-µm cell strainer, and cultured for expansion or immediate use.

#### Differentiation of human preadipocytes

As previously described (107), isolated SVC was expanded in DMEM/F12 containing FBS (10%) and P/S (1%), split to expand, and cultured until confluence. When cells reached 100% confluence, differentiation was induced using differentiation media, DM-2 (Zenbio, Inc) consisting of insulin, dexamethasone and isobutylmethylxanthine (IBMX) and pioglitazone supplemented with 10% FBS. After 3 days, the media was changed to adipocyte maintenance media, AM-1 (Zenbio, Inc) containing only insulin, dexamethasone in DMEM/F12 supplemented with 10% FBS (Fetal bovine serum). The cells were then left to differentiate for 3, 6, 12, and 14 days, with culture medium (AM-1) changed every 3 days. Day 0 preadipocytes were collected at time of confluence, just before adipogenesis induction. During differentiation, cells were collected at all the other time points and utilized immediately for experiments as described. Adipogenesis was confirmed by oil droplet formation and by increased expression of fatty acid-binding protein 4 gene, Fabp4.

#### SARS-CoV-2 infections of differentiated adipocytes, SVC, and A549-ACE2

Cells were seeded a day before infection by culturing 4×10^5^-1×10^6^ cells per well in a 6-well plate (Corning). On the day of infection, SVC was centrifuged at 500g for 5mins, and washed with infection media (DMEM, 2%FBS, and 1% Pen-strep). Adherent differentiated adipocytes and A549-ACE2 were washed with infection media. A549-ACE2 cells were cultured under the presence of 623µg/ml of Geneticin (Thermo Fisher Scientific; 10121035) for selection of ACE2 expressing cells. Viral infection was performed with SARS-CoV-2 (2019-nCOV/USA-WA1/2020) at a MOI of 1 for 1 hour while gently rocking before cells were washed and culture in culture media (DMEM, 10%FBS, and 1% Pen-strep) at 37°C with 5%CO_2_ under BSL3 containment. When remdesivir was used, cells were subjected to multiple washes with PBS 1 hour after infection before culturing under the presence of vehicle, dimethylsulfoxide (DMSO) (Sigma; D2650), or 10µM Remdesivir (Gilead) in DMSO and cultured for longer periods (24, 48, 72, and 96hpi).

#### SARS-CoV-2 infections of MA

MA media was changed into infection media by penetrating the fat tissue with a 22-gauge polytetrafluoroethylene (PTFE) (Grainger) blunt needle attached to a 3cc syringe (Grainger), or by gently transferring the fat with a wide manually cut p100 tip into a 5ml conical collection tube containing 2mls of warm media. Viral infection was performed with the WA/01 strain of SARS-COV-2 (2019-nCOV/USA-WA1/2020) at a multiplicity of infection (MOI) of 1 by gently adding virus to the top of the floating cells under BSL3 containment. MA was incubated for 1 hour while gently rocking at 37°C with 5%CO_2_. Media was then removed using as mentioned above prior to adding culture media (DMEM, 10%FBS, and 1% Pen-strep) and incubating at 37°C with 5%CO_2_.

#### Flow Cytometry

Cells were collected using Trypsin-EDTA (Thermo Scientific, 25200072) for 5-10mins at 37°C and 5% CO_2_. Cells were then centrifuged, 500g for 5 minutes, and washed with PBS (Thermo Scientific, 10010023) and transferred into a 96-well plate, then washed twice more with FACS buffer (PBS, 2%FBS (Corning, MT35016CV), and 5mM EDTA (Hoefer, GR123)). Cells were incubated in Zombie aqua cell viability dye (BioLegend, 423102) at room temperature for 20 minutes, washed twice, incubated at room temperature with Fc block (BioLegend, 422302) for 5 minutes, and surface stained for 30 minutes at 4°C. Surface staining contained antibodies for CD45 (BioLegend, 368503), CD3 (BioLegend, 317343), CD14 (BioLegend,301835), CD19 (BioLegend, 302261), CD11c (BioLegend, 371503), CD34 (BioLegend, 343615), CD31 (BioLegend, 303133). In fig. S4, surface staining also contained antibodies against ACE2 (R&D, fab9332r) or isotype control (R&D, ICOO3R). After surface staining, cells were washed twice with FACS buffer before fixation with 4% Paraformaldehyde (PFA) (Electron Microscopy Sciences, 15710) for 1 hour at 4°C, cells were then permeabilized (eBioscience, 00-8333-56) for 10 minutes at room temperature before intracellular staining for 45 minutes at 4°C. Intracellular staining contained an anti-SARS-CoV-2 N protein antibody (Sino Biological, 40143-T62). After intracellular staining, cells were washed with PBS, and diluted in 1%PFA and PBS before analyzing in a Cytek^TM^ Aurora. FCS files were collected and analyzed using FlowJo^TM^. Gating was done as shown in Fig. 1A.

#### 1- step RTqPCR for relative gene quantification

RNA was then mixed with TaqPath 1-step RTqPCR master mix (Applied Biosystems, A15299) and primers for SARS-CoV-2 *N-gene* (Biosearch technologies; n2019-nCoV KIT-NCOV-PP1-1000)*, IL-6* (Thermo scientific, Hs00174131_m1)*, IFNA1* (Thermo scientific, Hs04189288_g1)*, IFNB1* (Thermo scientific, Hs01077958_s1)*, ISG15* (Thermo scientific, Hs01921425_s1)*, IFI27* (Thermo Scientific, Hs01086373_g1)*,and IER3* (Thermo scientific, Hs00174674_m1) for relative gene quantification. Relative expression was calculated by obtaining ΔΔCT, using the endogenous control eukaryotic 18S rRNA (Thermo Scientific, 4352930E) and mock samples as the calibrator control. The samples were analyzed on a QuantStudio^TM^ 3 (Applied Biosystems) with the following parameters: 2 minutes at 25°C for uracil-DNA glycosylase (UNG) incubation, 15 minutes at 50°C for RT, 2 minutes at 95°C for polymerase activation, and 40 cycles of 15 seconds at 95°C and 30 seconds of 60°C for amplification.

#### 2- step RTqPCR for *ACE2, Fabp4,* and *18s*

RT on isolated RNA was performed with Superscript III first-strand synthesis system (Thermo scientific, 18080051). For qPCR, <100ng of cDNA were mixed with designed reverse and forward primers and PowerUp SYBR green master mix (Applied biosystems, A25742). The samples were analyzed on a QuantStudio^TM^ 3 (Applied Biosystems) with the following parameters: 10 minutes at 95°C for polymerase activation, and 40 cycles of 15 seconds at 95°C and 60 seconds of 60°C for amplification. The designed primers were the following: *ACE2 f*orward (TAACCACGAAGCCGAAGACC) and reverse (TTGGGCAAGTGTGGACTGTT), *Fabp4* forward (TGGGCCAGGAATTTGACGAA), and reverse (CACATGTACCAGGACACCCC); *18s* forward (GGCCCTGTAATTGGAATGAGTC) and reverse (CCAAGATCCAACTACGAGCTT). All primers were designed with mRNA sequence using National Center for Biotechnology Information primer designing tool, Primer-BLAST. All sequences are Sequence 5’→3’.

#### RNA*-*ISH

For deparaffinization, slides were baked at 70 °C for 1 hour in a temperature-controlled oven, then immersed in fresh xylene twice for 5 minutes each. Rehydration was performed using a Leica ST4020 Linear Stainer (Leica Biosystems) programmed to three dips per wash for 180 seconds each with the following buffers: xylene x 3, 100% ethanol x 2, 95% ethanol x 2, 80% ethanol x 1, 70% ethanol x 1, and ddH2O x 3. Heat induced epitope retrieval was subsequently performed in a Lab Vision PT module (Thermo Fisher) using the Dako Target Retrieval Solution, pH 9 (DAKO Agilent, S236784-2) at 97 °C for 10 minutes, followed by controlled cooling down to 65 °C. Slides were removed from the PT module and further cooled to room temperature for 30 minutes before rinsing briefly in sterile, nuclease-free water (ddH_2_O) twice. A 15 minutes hydrogen peroxide block was subsequently performed at 40 °C (Bio-Techne, 322335). Slides were then washed twice for 2 minutes each in ddH_2_O before an overnight hybridization at 40 °C with probes against the SARS-CoV-2 Spike mRNA (Bio-Techne, 848561). Amplification of the ISH probes was performed the next day according to the manufacturer’s protocol (Bio-Techne, 322350). Slides were then dried, mounted, and subsequently scanned via an Aperio slide scanner at the Stanford School of Medicine Histology Core.

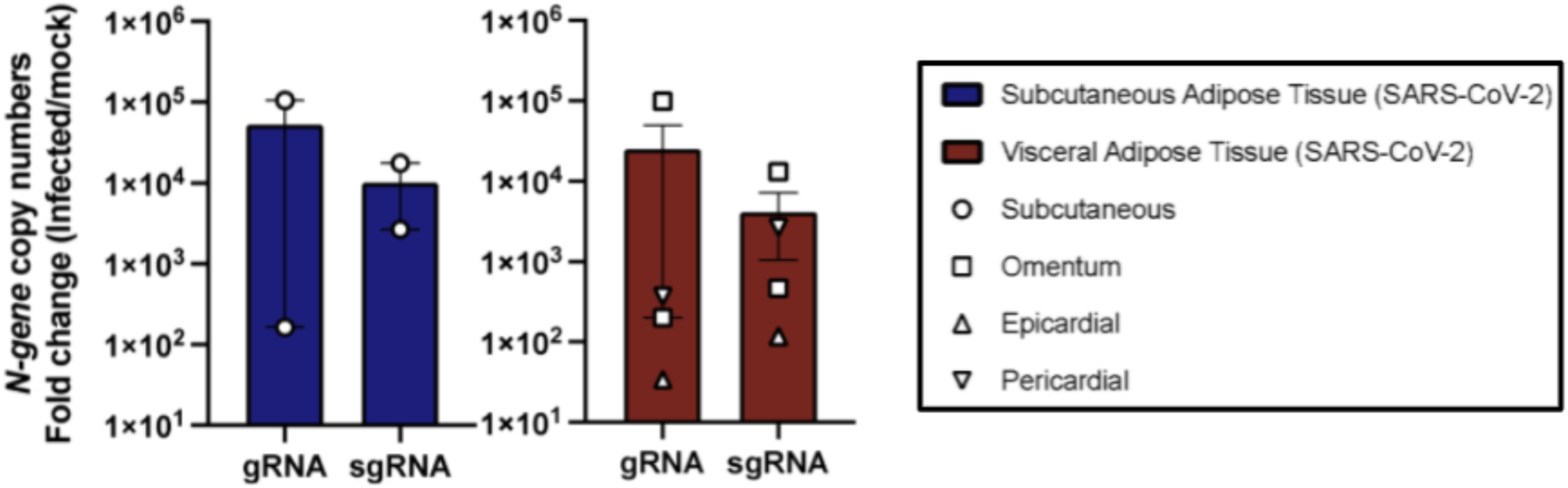

