## Supplemental Figures for "SARS-CoV-2 infects human adipose tissue and elicits an inflammatory response consistent with severe COVID-19"

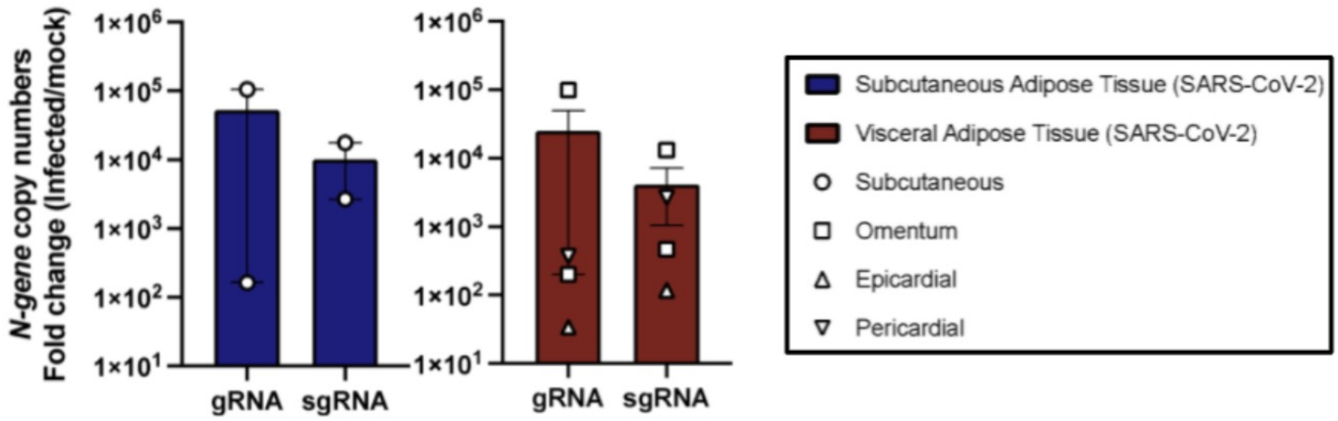

N-gene copy numbers
Fold change (Infected/mock)
1×10^6
1×10^5
1×10^4
1×10^3
1×10^2
1×10^1
gRNA
sgRNA
gRNA
sgRNA
Subcutaneous Adipose Tissue (SARS-CoV-2)
Visceral Adipose Tissue (SARS-CoV-2)
Subcutaneous
Omentum
Epicardial
Pericardial

**Supplementary Figure S1. Genomic and Subgenomic measurement of SARS-CoV-2 on *in vitro* infected SVC.** SVC was isolated by collagenase digestion from either SAT or VAT prior to viral infection. SVC was infected or left untreated (mock) for 24 hours with SARS-COV-2 (USA-WA1/2020) at a MOI of 1. RNA was obtained, and SARS-CoV-2 genome copy numbers infected subcutaneous (left; n=2) and visceral (right; n=4) adipose tissue was obtained by absolute gene quantification using 1-step RTQ-PCR TaqMan$^{TM}$ and reported as fold change of infected to mock. Data are presented as ± mean s.e.m.

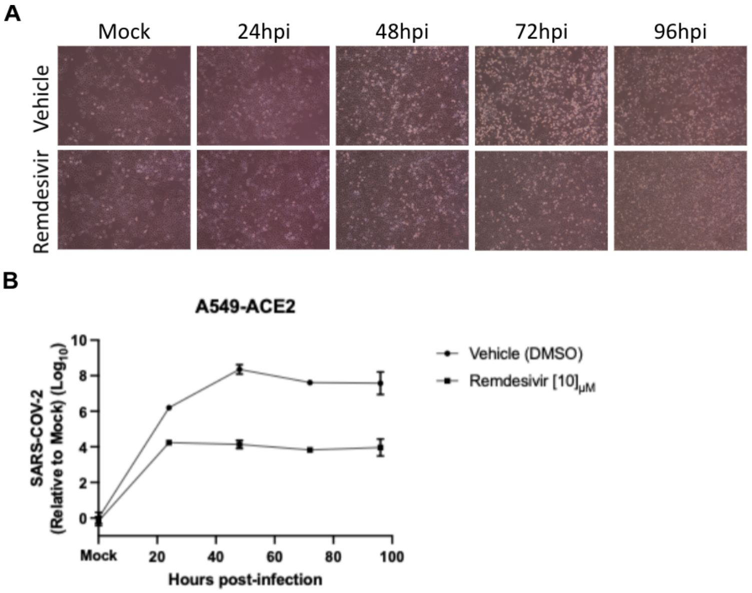


**Supplementary Figure S2. Validation of remdesivir treatments in A549-ACE2 cells.** A549-ACE2 cells were infected or left untreated (mock) for 1 hour with SARS-CoV-2 (USA-WA1/2020) at a MOI of 1, followed by washing and culturing in media containing vehicle (DMSO) or 10µM of remdesivir. Samples were harvested at 24, 48, 72, and 96 hpi. **(A)** cell culture images obtained with a EVOS XL Core Cell Imaging System showing cytopathic effect at 72hpi. **(B)** relative gene expression of SARS-CoV-2 (N gene) obtained by 1-step RTqPCR analyzed by ΔΔCt method relative to mock sample. Data represented in Log10. 18S rRNA was used as a housekeeping gene. Data are presented as ± mean s.e.m. Each point is an average of 2 biological replicates.

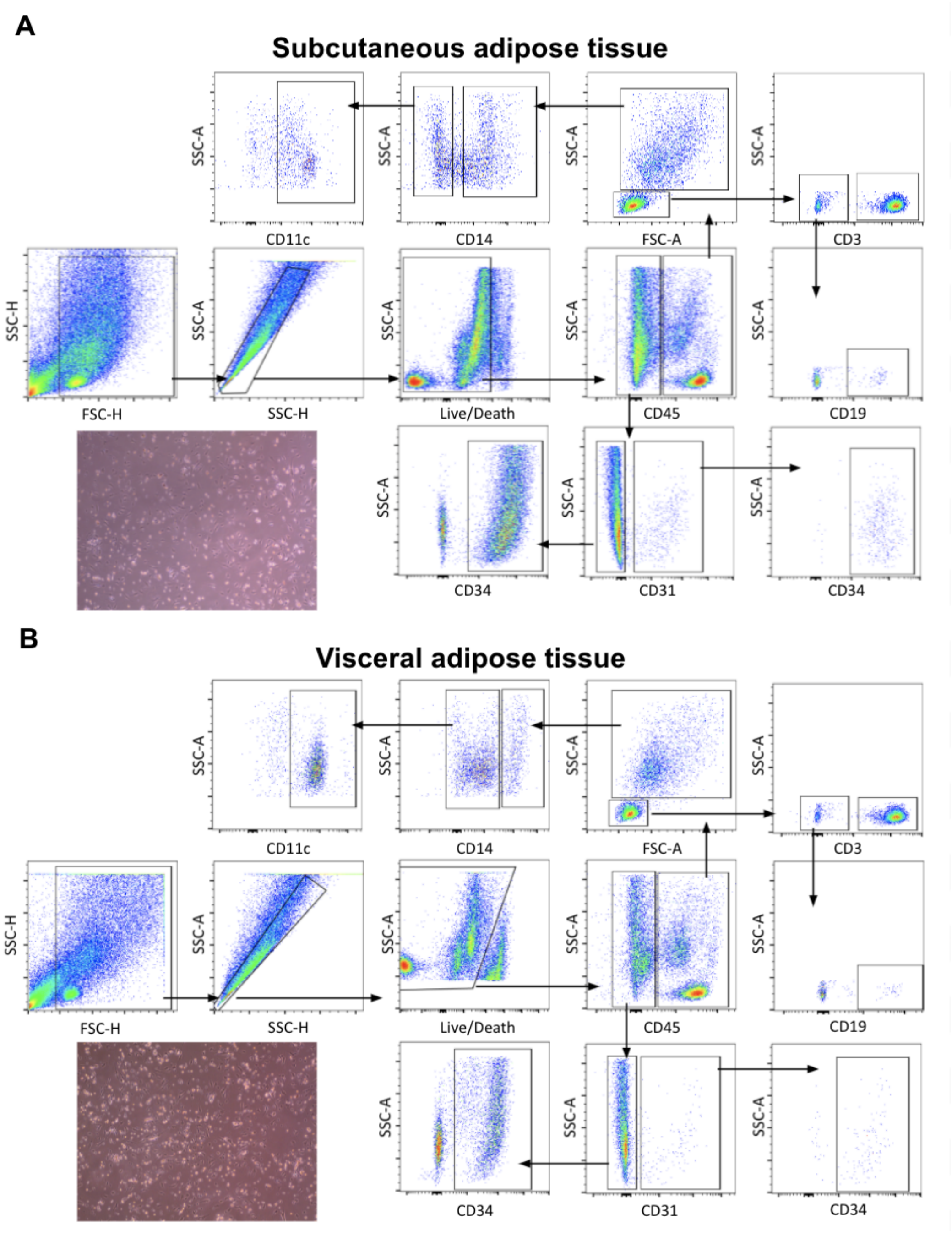

**Supplementary Figure S3. Gating strategy of SVC in human adipose tissue. (A)** Gating of SAT and **(B)** VAT SVC started by debris removal using forward (FSC) and side scatters (SSC) followed by removal of doublets by SSC-A and SSC-H and selection of viable cells by negative population of zombie aqua dye. Cells selection was done as followed: dendritic cells ($CD45^{+} \rightarrow$ myeloid $\rightarrow CD14^{-} \rightarrow CD11c^{+}$), macrophages ($CD45^{+} \rightarrow$ myeloid $\rightarrow CD14^{+}$), B cells ($CD45^{+} \rightarrow$ lymphocyte $\rightarrow CD3^{-} \rightarrow CD19^{+}$), T cells ($CD45^{+} \rightarrow$ lymphocyte $\rightarrow CD3^{+}$), endothelial cells ($CD45^{-} \rightarrow CD31^{+} \rightarrow CD34^{+}$), and preadipocytes ($CD45^{-} \rightarrow CD31^{-} \rightarrow CD34^{+}$). Flow cytometry data was collected using a 3-laser CYTEK Aurora; FCS files were analyzed in FlowJo$^{TM}$10. Cell culture images were obtained with a EVOS XL Core Cell Imaging System on healthy SVC.

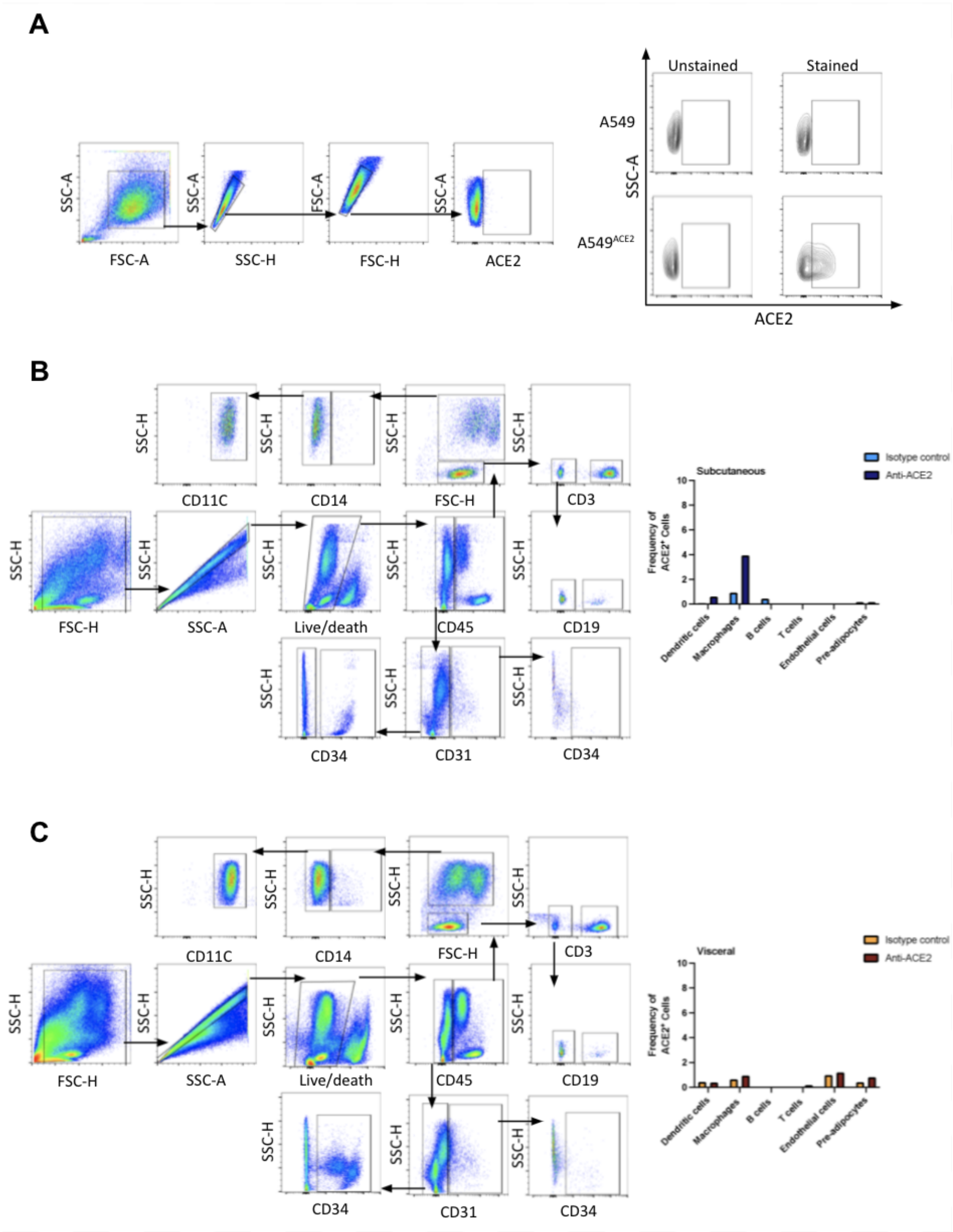


**Supplementary Figure S4. Limited ACE2 protein detection in SVC from SAT and VAT.** **(A)** Validation of commercially available human ACE2 antibody achieved by comparing cell lines deficient in ACE2 expression (A549) to genetically modified ACE2-expressing A549 cells, A549-ACE2. **(A)** To the left, gating strategy and to the right panel comparing unstained to stained A549 and A549-ACE2 cells. **(B-C)** Detecting ACE2 protein in SVC from SAT **(B)** and

VAT **(C)**; To the left, gating strategy as summarized in Fig.1A. To the right, results of frequency of ACE2 expression by comparing anti-ACE2 stained cells to isotype control.

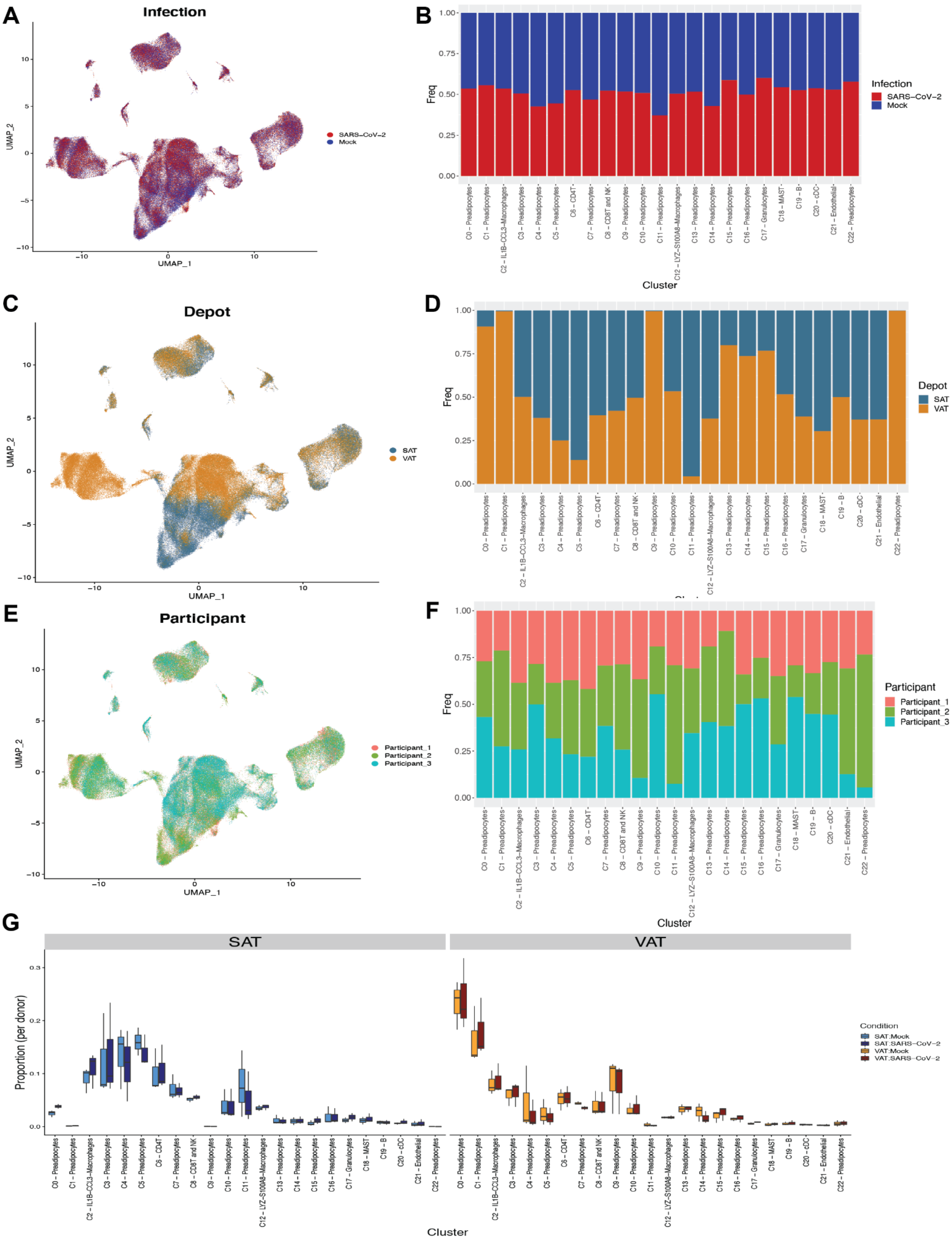
A
Infection
SARS–CoV–2
Mock
UMAP_1
UMAP_2
B
Freq
Cluster
Infection
C
Depot
SAT
VAT
D
Depot
E
Participant
Participant_1
Participant_2
Participant_3
F
Participant
G
SAT
VAT
Proportion (per donor)
Condition
SAT:Mock
SAT:SARS–CoV–2
VAT:Mock
VAT:SARS–CoV–2
C0 – Preadipocytes
C1 – Preadipocytes
C2 – IL1B–CCL3–Macrophages
C3 – Preadipocytes
C4 – Preadipocytes
C5 – Preadipocytes
C6 – CD4T
C7 – Preadipocytes
C8 – CD8T and NK
C9 – Preadipocytes
C10 – Preadipocytes
C11 – Preadipocytes
C12 – LYZ–S100A8–Macrophages
C13 – Preadipocytes
C14 – Preadipocytes
C15 – Preadipocytes
C16 – Preadipocytes
C17 – Granulocytes
C18 – MAST
C19 – B
C20 – cDC
C21 – Endothelial
C22 – Preadipocytes

**Supplementary Figure S5. Infection, depot and participant breakdown by cluster annotation.** (A, C, E) UMAP representation of SVC and (B, D, F) cell fraction barplot by cluster from all participants (n=3) across 198,759 cells, (A,B) colored by infection status, (C-D) colored by depot, and (E-F) colored by participant. (G) Barplot of the proportion of each cell type cluster within each participant by depot and infection status.

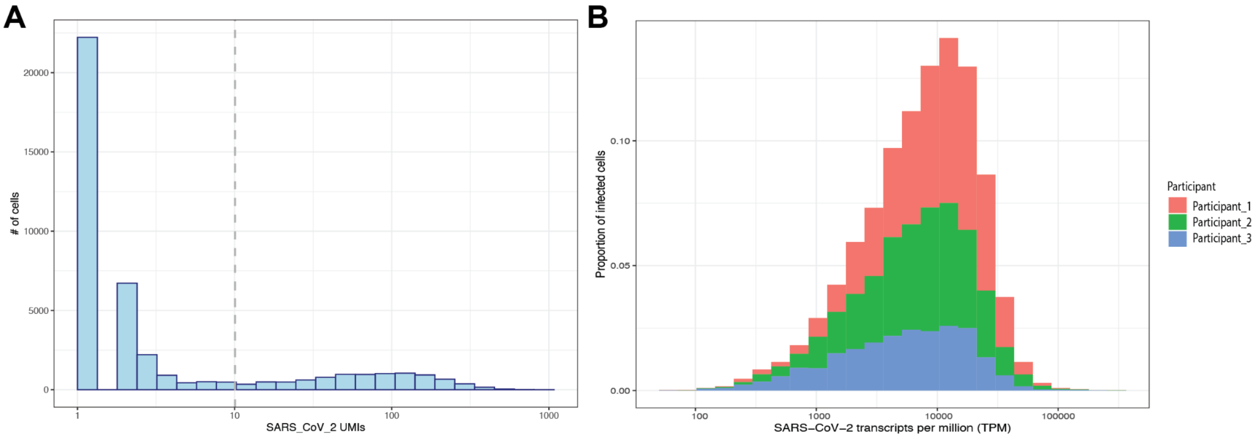


**Supplementary Figure S6. Distribution of SARS-CoV-2 transcripts across all cells.** (A) Distribution of SARS-CoV-2 UMIs across all cells with SARS-CoV-2 reads. The grey dotted line marks the 10UMI threshold used throughout. (B) Distribution of all cells with SARS-CoV-2 reads above threshold, colored by participant and plotted by SARS-CoV-2 cpm (counts per million).

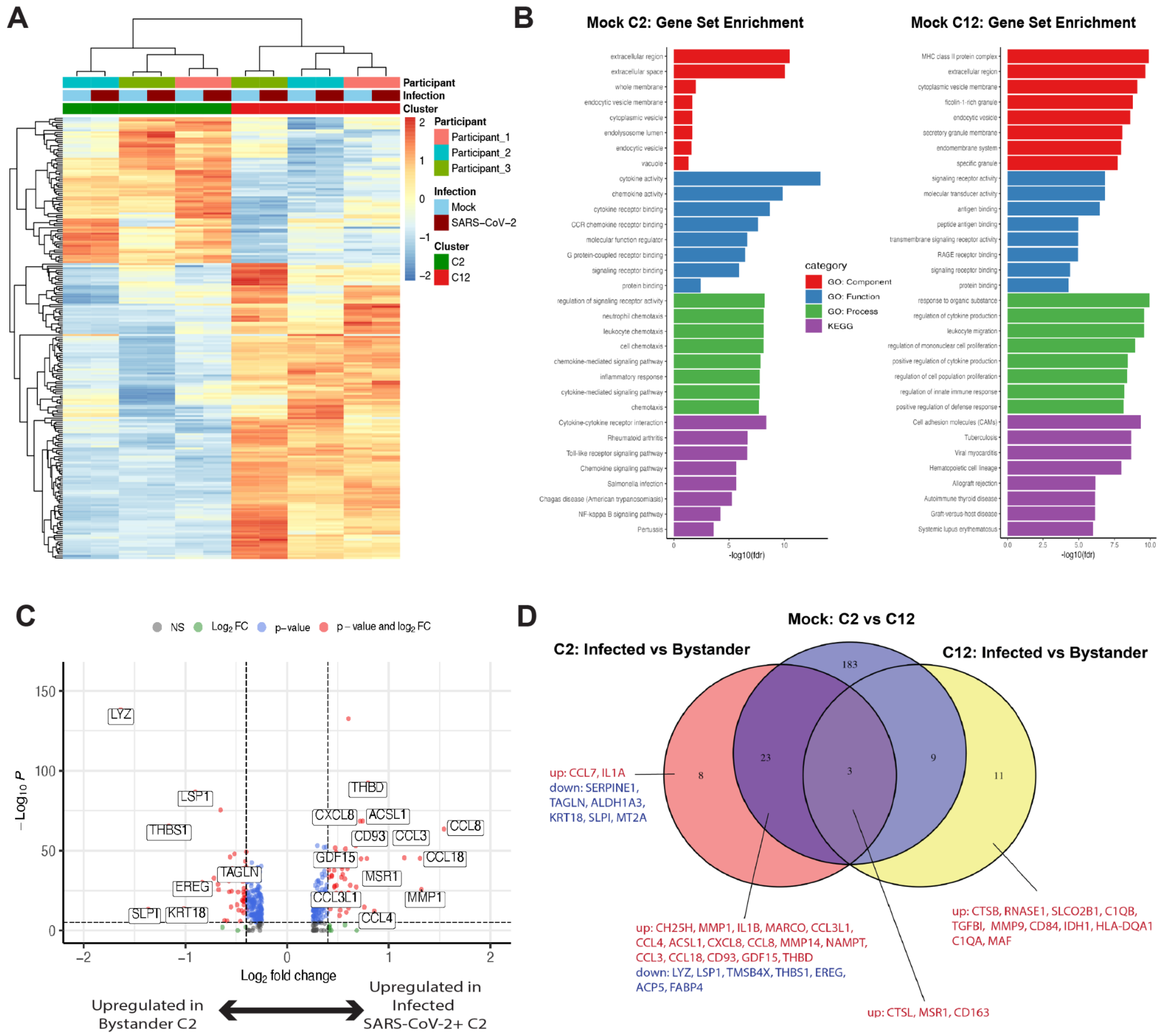


**Supplementary Figure S7. C2- and C12-macrophages are distinctly different upon mock and SARS-CoV-2 infection.** (A) Heatmap of the significant DEGs (padj < .01, abs(logfc) > 0.6) between mock C2- and C12-macrophages colored by z-score across all participants and infection conditions. (B) GO-term and KEGG pathway enrichment of the up-regulated genes derived from fig. S8A in C2 macrophages (74 DEGs) and C12 macrophages (146 DEGs). Top 10 pathways per pathway category with an FDR < 0.05 across each pathway category were included and the -log (FDR) was plotted per pathway. (C) Volcano plot of the differentially expressed genes between infected (SARS-CoV-2+) versus bystander C2-macrophages in SARS-CoV-2 infected

SAT and VAT. (D) Venn diagram comparing the significant DEGs (padj $< .01$, abs(logfc) $> 0.6$) with direction of change taken into account, across SARS-CoV-2+ infected macrophages versus bystander macrophages within the C2-macrophage population and within the C12-macrophage population of SARS-CoV-2 infected samples. DEGs between C2 and C12 across mock-infected samples were also included for comparison.

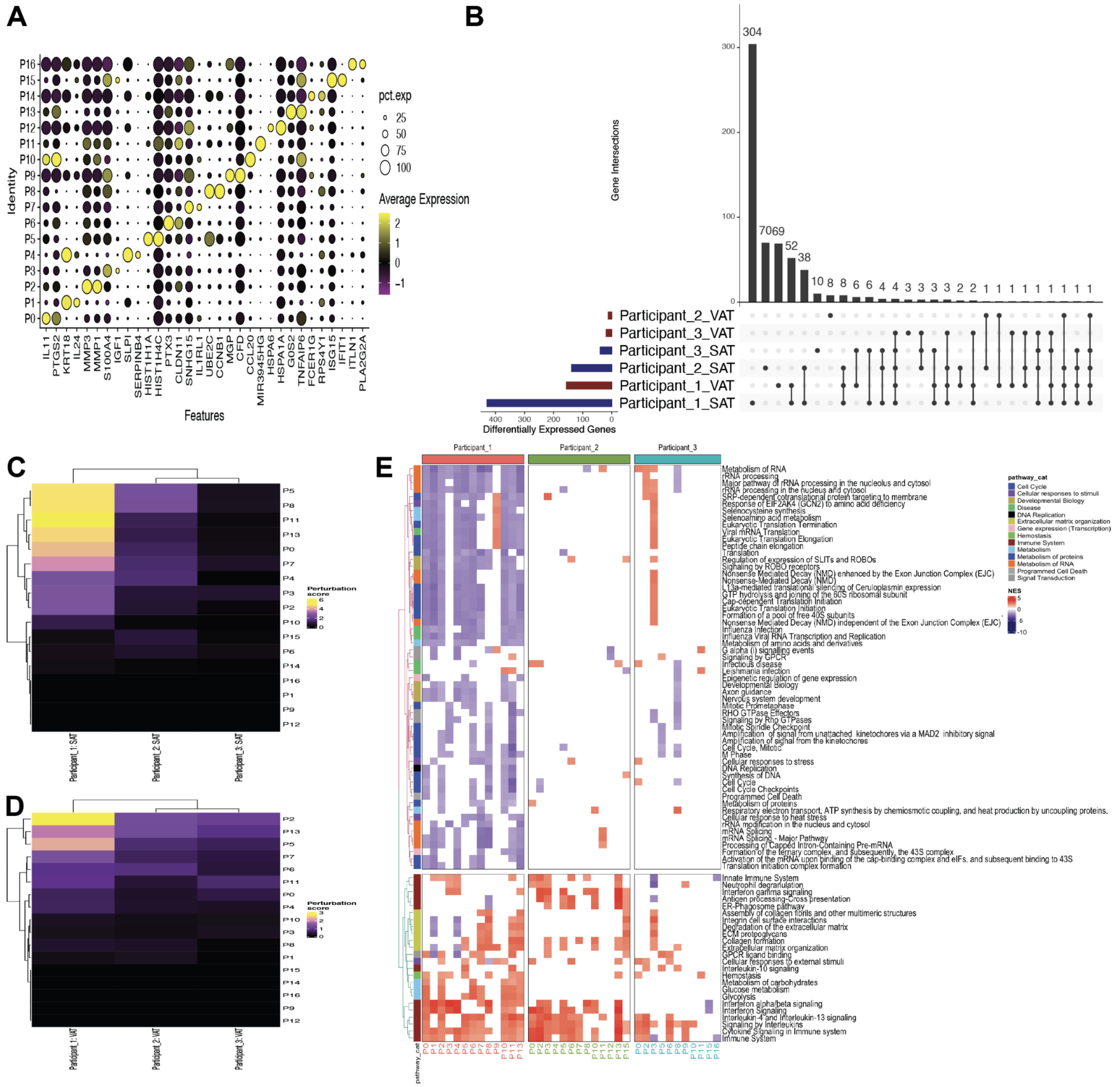


**Supplementary Figure S8. Characterization of preadipocytes across VAT and SAT.** (A) Dotplot of the top 2 cluster defining genes per preadipocyte cluster colored by scaled average expression. (B) Upset plot showing the differentially expressed genes in the preadipocytes of SARS-CoV-2 infected samples versus mock-infected samples. Left bar graph depicts the number of differentially expressed genes per depot and participant between the SARS-CoV-2 infected sample versus the mock-infected sample. The matrix depicts whether the DEGs are unique to the participant-depot or shared across multiple participant-depots. Bar graph at top demonstrates the number of DEGs that are associated with each participant-depot combination denoted by the matrix. (C, D) Perturbation analysis was performed by taking each preadipocyte cluster and first identifying the DEGs between SARS-CoV-2 infected samples versus the mock-infected samples and then projecting the whole transcriptome of each donor onto this gene set. Heatmaps of the perturbation scores per participant are displayed for both the(C) SAT and (D) VAT depots. (E) Reactome pathway analysis was performed on the significant DEGs by participants and clusters within VAT. Pathways that were represented and significant in at least four of the participant-cluster subsets were included. Pathways clustered by euclidean distance (tree not shown) and split by the two major subtrees.

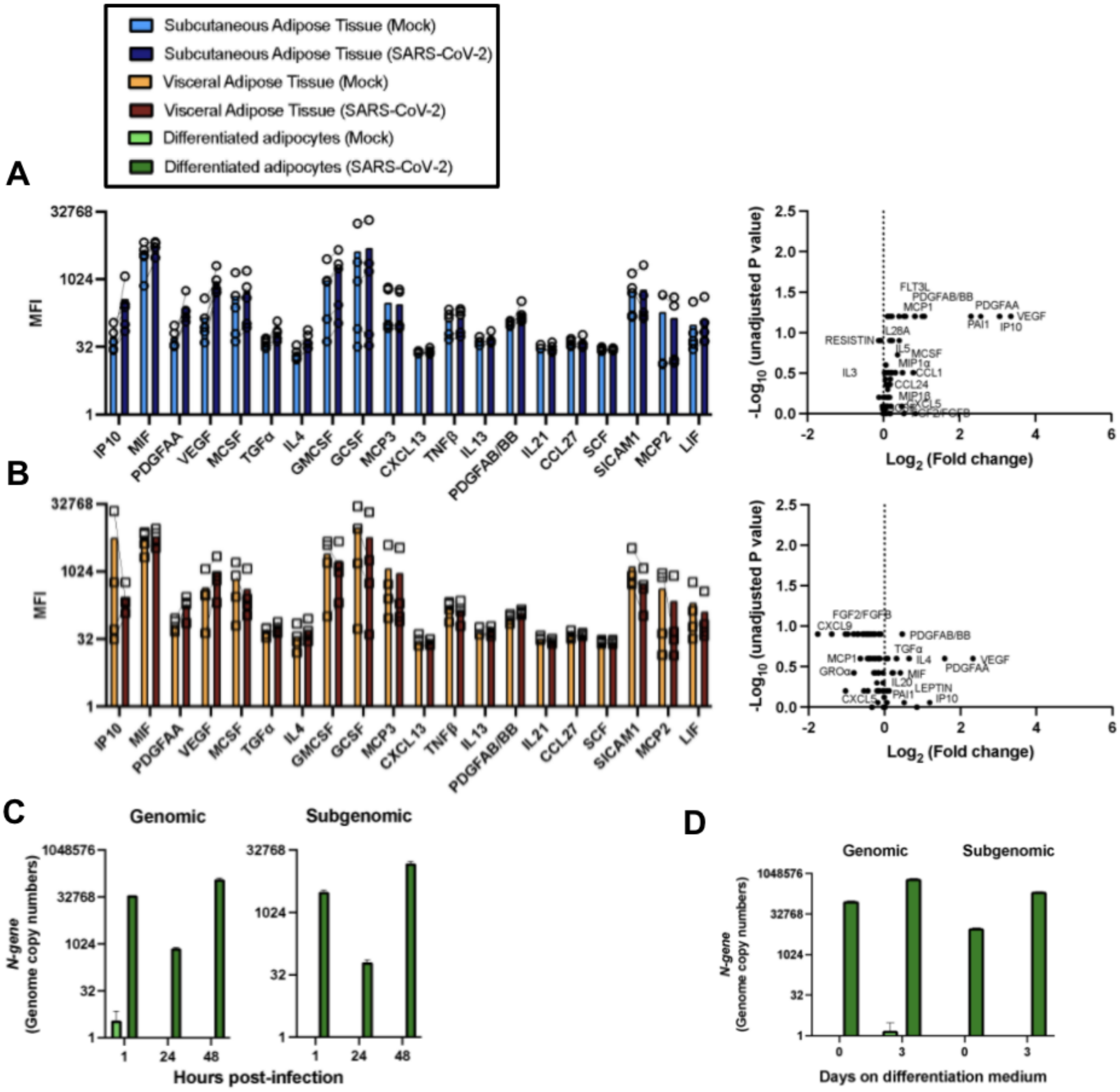


**Supplementary Figure S9. Mature and *in vitro* differentiated adipocytes harbor SARS-CoV-2 RNA and mount inflammatory responses after exposure to SARS-CoV-2 *in vitr*o.** Mature adipocytes (MA) of human adipose tissue were isolated by collagenase digestion prior to viral infection. MA were infected or left untreated (mock) for 24 hours with SARS-COV-2 (USA-WA1/2020) at a MOI:1. **(A-B)** Measurement of secreted analytes in the supernatant of mock and infected mature adipocytes from **(A)** SAT (n=5) and **(B)** VAT (n=4); left graph shows

20 of the 80 analytes analyzed by 80plex Luminex, right graphs are volcano plot with all 80 analytes. **(C-D)** Measurements of genomic and subgenomic SARS-CoV-2 genome copy numbers in mock and infected adipocytes differentiated from preadipocytes of subcutaneous adipose tissue with selection media for **(C)** 13 days (infected for 1, 24, and 48 hours) and **(D)** 0 or 3 days (infected for 24 hours); data was obtained by absolute gene quantification using 1-step RTqPCR TaqMan and reported as absolute values of genome copy numbers. Data presented as ± mean s.e.m.

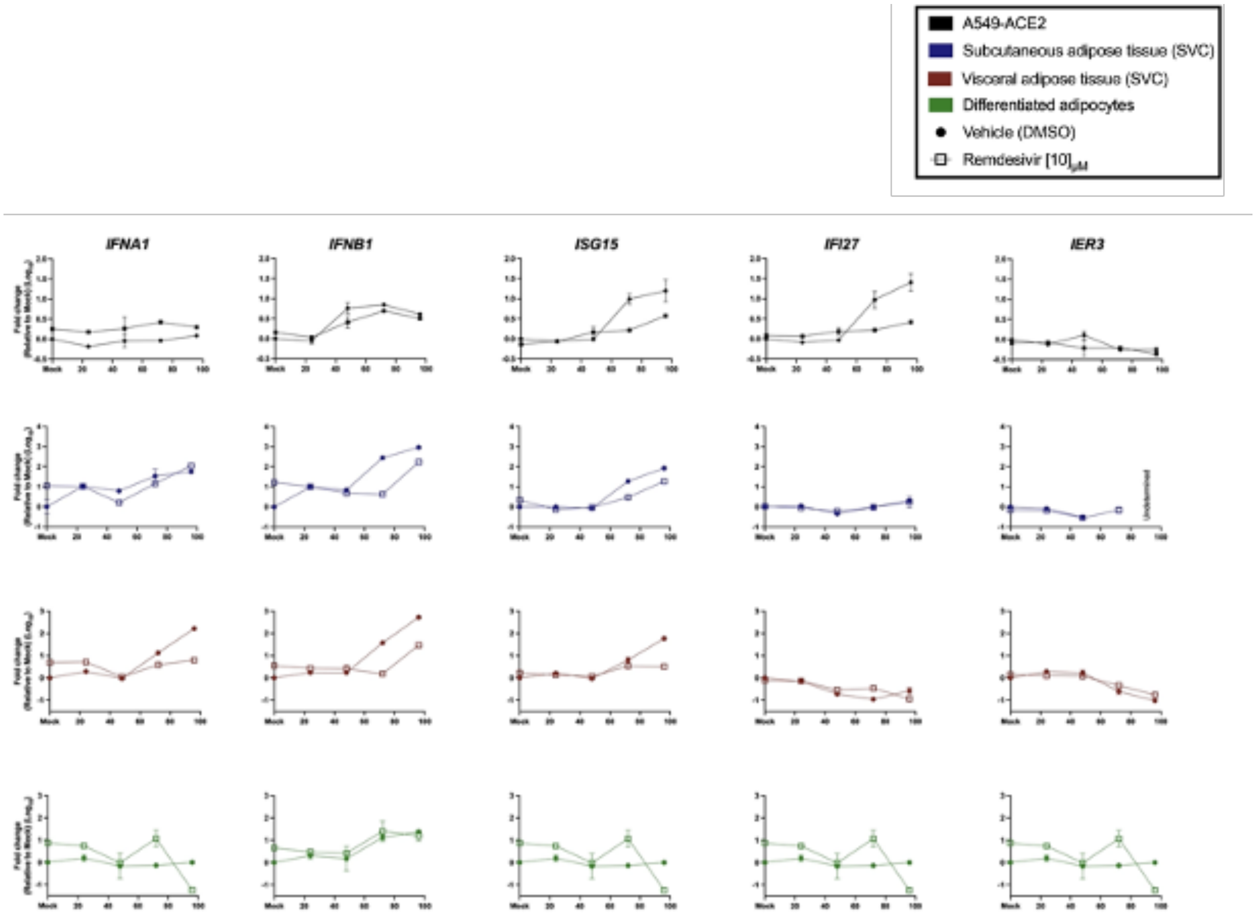


**Supplementary Figure S10. Increased interferon related genes in adipose tissue post-SARS-CoV-2 infection.** A549-ACE2 (Black), SAT-SVC (Blue), VAT-SVC (Red) and in vitro differentiated adipocytes (Green) were infected with SARS-CoV-2 (MOI:1) for 1 hours before washing the cells and culturing with either vehicle (DMSO) or remdesivir (10uM). RNA was isolated at 24, 48, 72, and 96 hours post infection. Relative quantification of *IFNA1, IFNB1, ISG15, IFI27, IER3* gene expression was obtained by 1-step RTqPCR using Taqman reagents. In all cases 18s was used as a housekeeping gene and data is shown as fold change (Log10) relative to mock (vehicle) sample.

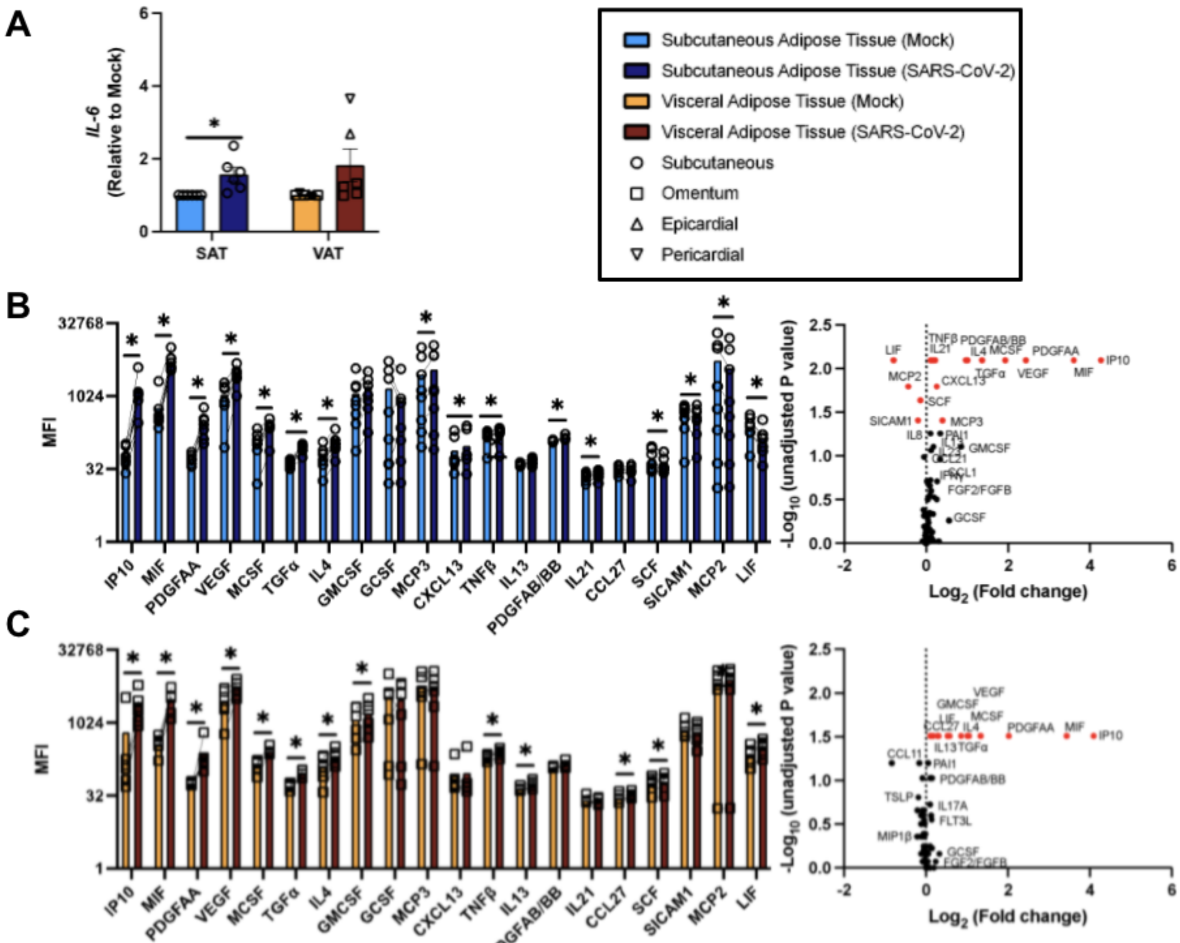


**Supplementary Figure S11. Elevated IL-6 gene expression and secretion of inflammatory mediators in SVC post in vitro SARS-CoV-2 infection.** Stromal vascular compartment (SVC) of human white adipose tissue was isolated by collagenase digestion prior to viral infection. SVC was infected or left untreated (mock) for 24 hours with SARS-CoV-2 (USA-WA1/2020) at a multiplicity of infection (MOI) of 1. **(A)** Relative gene expression of IL-6 (SAT, n=6; VAT, n=6) obtained by RTqPCR using eukaryotic 18S rRNA as a housekeeping gene and analyzed by the ΔΔCt method relative to mock samples. **(B)** Measurement of secreted analytes in the supernatant of mock and infected SVC from SAT (Top, n=8) and **(C)** VAT (bottom, n=6). The left graph shows 20 of the 80 analytes analyzed by 80-plex Luminex, right graphs are volcano plot with all 80 analytes. Statistical analysis: **(A)** paired, two-sided, student's t-test. *$P<0.05$ **(E-F)** paired Wilcoxon signed rank test; *analytes that were significantly different with unadjusted P values. Data are presented as ± mean s.e.m. Red dots in **(B-C)** volcano plot represents analytes that significantly differed from mock with $P<0.05$.

**Table S1: Adipose tissue donor's demographic, medical information, and sample use**

| ID | Age | Sex | Ethnicity | T2DM/ HTN | BMI | Medications | Other risk factors | Adipose depot | Experiment performed |
|---|---|---|---|---|---|---|---|---|---|
| 531 | 59 | M | W | Y/Y | 41 | Metformin, Lantus, Ozempic, Jardiance, Atenolol, and Lisinopril | Severe obesity | SAT, and VAT | RTqPCR (ACE2, SARS-CoV-2), RTqPCR (ACE2) on differentiated adipocytes |
| 888 | 49 | F | W | Y/Y | 52 | Dulaglutide, Empagliflozin, Metformin, Carvedilol, Anastrozole, and Lisinopril | Severe obesity, CHF, and breast Cancer | SAT, and VAT | Used to develop experimental protocols |
| 108 | 36 | F | H | N/Y | 47 | Amlodipine, and Lisinopril and Hydrochlorothiazide | Severe obesity, and Asthma | SAT and VAT | Used to develop experimental protocols |
| 935 | 58 | F | W | N/N | 37 | None | Obesity | SAT and VAT | RTqPCR (ACE2) on differentiated adipocytes |

| | | | | | | | | | |
|---|---|---|---|---|---|---|---|---|---|
| 159 | 52 | M | H | N/N | 40 | Liraglutide, and Metformin | Severe obesity | SAT and VAT | Used to test antibodies for Flow cytometry panel |
| 120 | 33 | F | H | N/N | 52 | None | Severe obesity | SAT and VAT | Luminex, RTqPCR (ACE2, IL-6, SARS-CoV-2) |
| 611 | 52 | F | W | N/Y | 46 | Valsartan, and Hydrochlorothiazide | Severe obesity | SAT and VAT | Luminex, Flow cytometry (SARS-CoV-2) |
| 959 | 54 | F | W | N/Y | 40 | Valsartan, and Hydrochlorothiazide | Severe obesity | SAT and VAT | Luminex |
| 608 | 42 | M | W | N/Y | 52 | Metformin, and Liraglutide | Severe obesity | SAT and VAT | Luminex, Flow cytometry (SARS-CoV-2) |
| 002 | 67 | F | W | Y/Y | 30.1 | Glyburide, Metformin, Hydrochlorothiazide, and Lopressor | Obesity, Asthma, and CAD | EAT, PAT, and SAT | Used to develop experimental protocols |
| 003 | 61 | M | H | N/Y | 30.8 | Labetalol, Valsartan, and Amlodipine | Obesity, CHF, and CAD | EAT, PAT, and SAT | RTqPCR (SARS-CoV-2) on differentiated adipocytes |
| 004 | 72 | F | H | Y/Y | 33.2 | Amlodipine, and Lopressor | Obesity, ESRD on HD, and CAD | EAT, PAT, and SAT | RTqPCR (ACE2) on differentiated adipocytes and remdesivir timecourse |

| | | | | | | | | | |
|---|---|---|---|---|---|---|---|---|---|
| 005 | 63 | M | H | Y/Y | 29.4 | Canagliflozin, Glipizide, Insulin glargine, Metformin, Benazepril, Furosemide, and Lopressor | Smoker, and CAD | EAT, PAT, and SAT | Luminex, RTqPCR (ACE2, IL-6, SARS-CoV-2) |
| 319 | 61 | F | W | N/N | 32.8 | None | Obesity | SAT | Luminex |
| 209 | 35 | F | H | N/N | 38.4 | None | Obesity | SAT and VAT | Flow cytometry (ACE2) |
| 272 | 36 | F | A | N/Y | 42.8 | Losartan | Severe obesity | SAT and VAT | Luminex, and remdesivir timecourse |
| 658 | 60 | F | H | Y/N | 38.1 | Metformin, and Azathioprine | COPD, Asthma, Obesity, Smoker, and MG | SAT and VAT | scRNA-seq, Luminex, and RTqPCR (SARS-CoV-2, and IL-6) |
| 576 | 33 | M | H | Y/Y | 47.9 | Metformin | Severe obesity | SAT and VAT | scRNA-seq, Luminex, and RTqPCR (SARS-CoV-2, and IL-6) |
| 924 | 50 | F | B | N/Y | 54.5 | Insulin glargine, Insulin lispro, Losartan, Furosemide, and Carvedilol | T1DM, severe obesity, former smoker | SAT and VAT | scRNA-seq, Luminex, and RTqPCR (SARS-CoV-2, and IL-6) |

**Abbreviations:** subcutaneous adipose tissue (SAT), visceral adipose tissue (VAT), epicardial adipose tissue (EAT), pericardial adipose tissue (PAT), reverse transcription quantitative polymerase chain reaction (RTqPCR), Single cell ribonucleic acid sequencing (scRNA-seq), type 1 diabetes mellitus (T1DM), type 2 diabetes mellitus (T2DM), hypertension (HTN), male (M), female (F), yes (Y), no (N), body mass index (BMI), white (W), black (B), hispanic (H), asian (A), congestive heart failure (CHF), coronary artery disease (CAD), end-stage renal disease (ESRD), hemodialysis (HD), and chronic obstructive pulmonary disease (COPD), myasthenia gravis (MG).

**Supplementary Table S1. Adipose tissue participant's demographic, medical information, and sample use.** Table summarizing all adipose tissue participant information from samples used in this study. The information presented is the following: (left to right) participant ID, age, sex, ethnicity, if patient had type 2 diabetes and/or hypertension, body mass index (BMI), other risk factors, type of adipose tissue obtained, and experiments performed on adipose tissue.

**A** ***ACE2* Ct values from adipose tissue**

| | | Ct value | |
|---|---|---|---|
| **Subject ID** | **Adipose tissue** | **Stromal vascular compartment** | **Mature adipocytes** |
| 005 | Subcutaneous | NS | NS |
| | Epicardial | NS | 37 |
| | Pericardial | NS | NS |
| 120 | Subcutaneous | NS | NS |
| | Visceral | NS | NS |
| 531 | Visceral | NS | NS |

**Note:** RNA from human kidney was used as a positive control for *ACE2* primers (Ct=30). All Ct values ≥ 37 were considered as no signal (NS).

**B** **Ct values of *ACE2*, *Fabp4*, and *18s* on preadipocytes in differentiation media**

| | | | Ct | | |
|---|---|---|---|---|---|
| Subject ID | Adipose tissue | Days in differentiation media | *ACE2* | *Fabp4* | *18s* |
| 935 | Subcutaneous | 0 | No signal | 35.1 | 14.8 |
| | | 3 | 35.0 | 24 | 12.5 |
| | | 6 | No signal | 19.8 | 13.1 |
| | | 14 | 35.9 | 22.8 | 14.2 |
| 532 | Visceral | 0 | 37 | 37 | 14.9 |
| | | 3 | 34 | 29.6 | 14.8 |
| | | 6 | 35.5 | 27.7 | 14.6 |
| | | 14 | 35 | 27.6 | 14.2 |
| 004 | Paracardial | 0 | No signal | 36.2 | 14.1 |
| | | 3 | 33.4 | 26.3 | 14.1 |
| | | 6 | 34.5 | 27 | 15.1 |
| | | 13 | 36.9 | 27.7 | 14.1 |

Note: Ct values are an average of two technical replicates.

**Supplementary Table S2. No expression of *ACE2* in adipose tissue but increased ACE2 expression in *in vitro* differentiated adipocytes at 3 days of differentiation. (A)** Table summarizing *ACE2* Ct values from SVC and mature adipocytes of adipose tissue obtained from subcutaneous, visceral, epicardial and pericardial fat from patients 005, 120, and 531. **(B)** Table summarizing *ACE2*, *Fabp4*, and *18s* Ct values from preadipocytes on differentiation media for 0, 3, 6, 13 or 14 days. SVC for differentiation of preadipocytes was obtained from either SAT, VAT, or PAT from patients 935, 532, and 004. (A-B) Ct values were obtained by 2-step RTqPCR.
